# Male meiotic spindle features that efficiently segregate paired and lagging chromosomes

**DOI:** 10.1101/737494

**Authors:** Gunar Fabig, Robert Kiewisz, Norbert Lindow, James A. Powers, Vanessa Cota, Leslie Mateo, Jan Brugués, Steffen Prohaska, Diana S. Chu, Thomas Müller-Reichert

## Abstract

Chromosome segregation during male meiosis is tailored to rapidly generate multitudes of sperm. Little, however, is known about the mechanisms that efficiently segregate chromosomes to produce sperm. Using live imaging in *Caenorhabditis elegans*, we find that spermatocytes exhibit simultaneous pole-to-chromosome shortening (anaphase A) and pole-to-pole elongation (anaphase B). Electron tomography unexpectedly revealed that spermatocyte anaphase A does not stem from kinetochore microtubule shortening. Instead, movement is driven by changes in distance between chromosomes, microtubules, and centrosomes upon tension release at anaphase onset. We also find that the lagging X chromosome, a distinctive feature of anaphase I in *C. elegans* males, is due to lack of chromosome pairing. The unpaired chromosome remains tethered to centrosomes by continuously lengthening kinetochore microtubules which are under tension, suggesting a ‘tug of war’ that can reliably resolve chromosome lagging. Overall, we define features that partition both paired and lagging chromosomes for optimal sperm production.

## Introduction

Chromosome segregation during meiosis is regulated in each sex to produce different numbers of cells of distinct size, shape, and function. In humans, for example, up to 1500 sperm are continually generated per second via two rapid rounds of symmetric meiotic divisions. In contrast, only oocytes that are fertilized will complete asymmetric meiotic divisions to produce one large cell and two to three small polar bodies (El Yakoubi and Wassmann, 2017; L’Hernault, 2006; O’Donnell and O’Bryan, 2014; Severson et al., 2016; Shakes et al., 2009). While oocyte meiosis and also mitosis have been studied in detail in many organisms (Bennabi et al., 2016; Muller-Reichert et al., 2010; Pintard and Bowerman, 2019), our current knowledge of sperm meiotic chromosome segregation is limited to studies using chromosome manipulation and laser microsurgery in grasshoppers and crane flies (LaFountain et al., 2011; LaFountain et al., 2012; Nicklas and Kubai, 1985; Nicklas et al., 2001; Zhang and Nicklas, 1995). Thus, despite recent evidence of steep global declines in human sperm counts (Levine et al., 2017; Levine et al., 2018; Sengupta et al., 2018), little is known about the fundamental molecular mechanisms of male meiotic chromosome segregation that are required for efficiently forming healthy sperm.

Spermatocytes exhibit vastly distinct spindle structures and dynamics with respect to spindle poles, chromosomes, and kinetochores compared to oocytes or cells in mitosis (Crowder et al., 2015; Hauf and Watanabe, 2004). In many species, centrosomes are present in spermatocyte meiosis and mitosis but not in oocyte meiosis, which highly influences microtubule organization that specifies differences in spindle structure, size, and shape. Chromosomes also adopt different shapes, potentially altering the types and degree of interactions with microtubules. In the model organism *Caenorhabditis elegans*, for example, chromosomes resemble compact oblong spheres in spermatocyte and oocyte meiosis (Albertson and Thomson, 1993; Redemann et al., 2018; Shakes et al., 2009) but are long rods in mitosis (Oegema et al., 2001; Redemann et al., 2017). Kinetochore structure and dynamics during meiosis also vary from mitosis, where kinetochores attach each sister chromatid to microtubules that distribute sisters to opposite poles during the single division. During meiosis I when homologs separate, kinetochores on sister chromatids attach to microtubules from the same pole. Kinetochores must then somehow detach and re-attach to microtubules to allow sisters to then segregate to opposite poles for meiosis II (Petronczki et al., 2003). In acentrosomal oocyte meiosis, this is accomplished because outer kinetochore levels dramatically decrease during anaphase I and then increase in preparation for meiosis II (Dumont and Desai, 2012; Dumont et al., 2010). The structure and dynamics of centrosomal spermatocyte meiotic spindles that achieve efficient meiotic chromosome segregation, however, are largely unknown.

Distinct spindle structures can also drive chromosome movement by different means (McIntosh, 2017; McIntosh et al., 2012). During anaphase A, chromosome-to-pole distance decreases, while pole-to-pole distance is constant (Asbury, 2017). Typically, a shortening of kinetochore microtubules drives this anaphase A movement (Asbury, 2017). During anaphase B, the poles move closer to the cortex, causing the pole-to-pole distance to increase while chromosome-to-pole distance is constant (Scholey et al., 2016). Forces that separate poles are the pulling force acting on astral microtubules and the pushing force from microtubules forming in the spindle midzone as chromosomes separate. Systems differ in use of these means. *C. elegans* uses solely anaphase B during the first mitotic division (Nahaboo et al., 2015; Oegema et al., 2001). In *C. elegans* oocyte spindles, a shortening of the distance between chromosomes and the acentrosomal poles was observed before microtubules disassemble at the poles (McNally et al., 2016). Pushing forces generated by microtubules that assemble in the spindle midzone are suggested to drive the majority of segregation in the oocyte (Laband et al., 2017; Yu et al., 2019). Thus depending upon cellular context, different spindle structures will balance mechanisms and forces to drive chromosome movement. As yet, the mechanisms that drive segregation in sperm centrosomal meiotic spindles are unknown.

*C. elegans* is ideal to study chromosome segregation in spermatocytes. Sperm can be visualized by both light and electron microscopy in both sexes. *C. elegans,* which lacks a Y chromosome, determines sex by X chromosome number: hermaphrodites have two X chromosomes (XX), while males have one X (XO). Light microscopy has shown that during meiosis I, the unpaired (univalent) X chromosome in XO males lags during anaphase I (Albertson and Thomson, 1993; Fabig et al., 2016; Madl and Herman, 1979). Electron microscopy has defined the ultrastructural organization of *C. elegans* spindles in meiotic oocytes (Laband et al., 2017; Redemann et al., 2018; Srayko et al., 2006; Yu et al., 2019) and mitotic embryos (Albertson, 1984; O’Toole et al., 2003; Redemann et al., 2017; Yu et al., 2019) and unrevealed the holocentric nature of the *C. elegans* kinetochore (Albertson and Thomson, 1993; Howe et al., 2001; O’Toole et al., 2003). However, ultrastructural studies on chromosome and spindle organization during meiotic segregation have not been performed for spermatocytes, hindering our understanding of sex-specific regulation of meiotic chromosome segregation.

Given that sperm meiosis has both similarities and differences to oocyte meiosis and mitosis, a key question applicable to a broad range of organisms is how spindle structure and dynamics are regulated in sperm to navigate two rounds of centrosomal meiotic divisions, including the efficient resolution of lagging chromosomes. To answer this, we applied a combination of light microscopy with a newly developed approach for imaging meiosis in whole living males, high-resolution immunostaining, and electron tomography, which produces a large-scale 3D reconstruction of spermatocyte meiotic spindle structure. The combination of these results defines molecular mechanisms of sperm-specific movements that are driven by interactions of microtubules with chromosomes and spindle poles as they navigate two rounds of centrosomal meiotic divisions.

## Results

### Spermatocyte meiotic spindles are distinguished by delayed segregation of the unpaired X chromosome

To visualize chromosome dynamics in spermatocyte meiosis, we developed *in situ* imaging of spermatocytes within *C. elegans* males to visualize microtubules and chromosomes labeled with β-tubulin::GFP and histone::mCherry, respectively (**Fig. 1**). The chromosomes arrange in a rosette pattern, with paired autosomes surrounding the unpaired X chromosome at metaphase I (Albertson and Thomson, 1993). At anaphase I, homologs segregate towards opposite poles; the unpaired X chromosome, however, is left behind, attached to microtubules that connect to separating poles (**Fig. 1A**, arrowheads) (Albertson and Thomson, 1993). Later in anaphase I, the X moves towards one of the spindle poles (**Suppl. Movie S1**). Thus, it appears that sister chromatids of the unpaired X attach to opposite poles in meiosis I in contrast to the autosomes, in which sister chromatids attach to the same pole to enable segregation of homologs to opposite poles. Meiosis I, therefore, shows differences in the segregation of paired chromosomes compared to the unpaired X, in contrast to the second division, where sister chromatids of all chromosomes segregate to opposite poles (**Fig. 1B, Suppl. Movie S1**).

**Figure 1.**
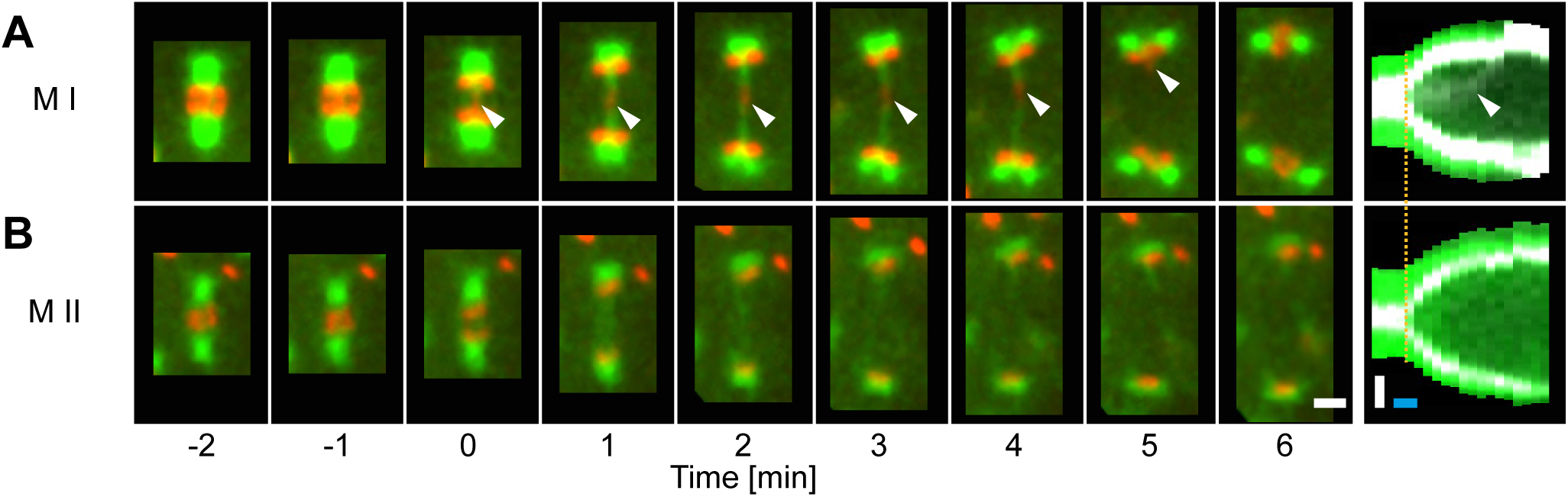
Spindle and X chromosome dynamics during meiotic divisions in males. Time series of confocal image projections of (**A**) meiosis I (M I) and (**B**) meiosis II (M II). Microtubules (β-tubulin::GFP, green) and chromosomes (histone H2B::mCherry, red) are visualized. Anaphase onset is time point zero (t=0). The progression of chromosome segregation is visualized in kymographs (right panels; anaphase onset is indicated by an orange line). The position of the unpaired X chromosome in meiosis I is marked by white arrowheads. Scale bar (white), 2 µm; time bar (blue), 2 min.

### Lagging of chromosomes is a consequence of a lack of pairing

We probed if the lagging of the X may be due to a lack of having a pairing partner. Because both males and hermaphrodites undergo spermatogenesis in *C. elegans*, we compared the behavior of chromosomes in spermatocytes of wild-type males (XO) to those in animals with different numbers of chromosomes. First, though the unpaired X chromosome lags in wild-type male (XO) spermatocytes (**Fig. 2A**), paired X chromosomes in wild-type hermaphrodite (XX) spermatocytes did not (**Fig. 2B**). Further, we determine whether paired X chromosomes lag in males by analyzing mutant males that have two sex chromosomes (XX). For this, we imaged animals with the *tra-2*(e1094) mutation, which causes somatic transformation of XX animals to males (Hodgkin and Brenner, 1977). In over 80% (n = 43/53) of *tra-2*(e1094) spindles we did not detect lagging chromosomes during meiosis I (**Fig. 2C**). Therefore, paired X chromosomes in male spermatocyte meiosis do not lag, similar to paired sex chromosomes in hermaphrodites.

**Figure 2.**
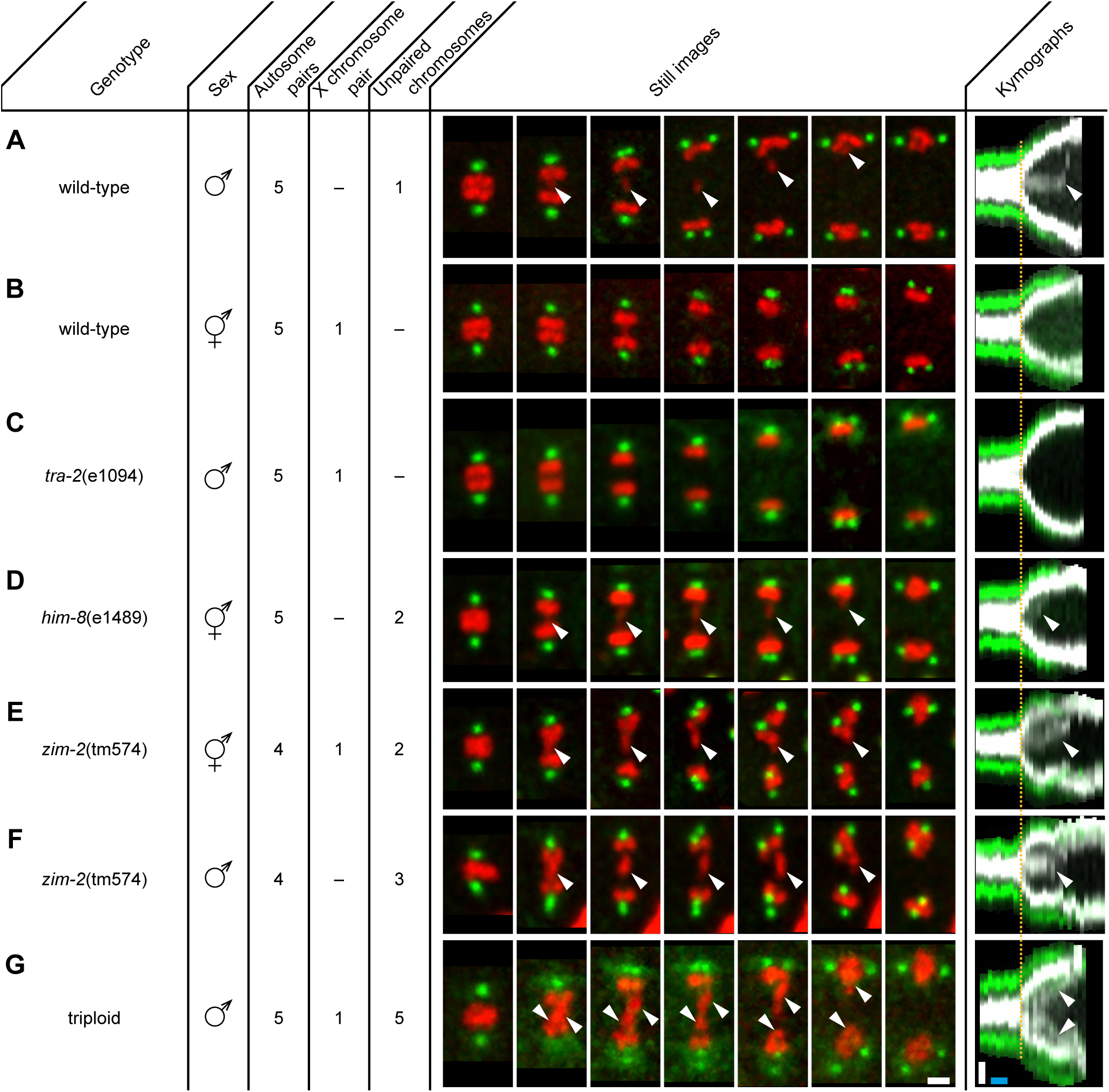
Unpaired chromosomes lag in male spermatocyte meiosis I. Confocal image projections of spermatocyte meiosis I in (**A**) wild-type XO males, (**B**) wild-type XX hermaphrodites, (**C**) *tra-2*(e1094) XX males, (**D**) *him-8*(e1489) XX hermaphrodites, (**E**) in *zim-2*(tm574) XX hermaphrodites, (**F**) *zim-2*(tm574) XO males, (**G**) in triploid XXO males. Centrosomes are labeled in green (γ-tubulin::GFP) and chromosomes in red (histone H2B::mCherry). The genotype, sex, number of autosome pairs, occurrence of paired X chromosomes and number of unpaired chromosomes is indicated. Still images illustrate the progression of the first meiotic division over time. Arrowheads (white) indicate lagging chromosomes. In the corresponding kymographs (right panels), chromosomes are shown in gray, spindle poles in green. Anaphase onset is marked (dashed line, orange). Scale bars (white), 2 µm; time bars (blue), 2 min.

We next examined X chromosome lagging in *him-8*(e1489) hermaphrodites. Observing lagging in hermaphrodites would eliminate the possible effect of the male soma causing chromosomes to lag in meiosis I. Further, mutation of *him-8* results in lack of recognition of the pairing center on the X chromosome; thus, pairing, synapsis, and recombination of the X chromosomes do not occur (Phillips et al., 2005). In 70% (n = 14/20) of the analyzed *him-8*(e1489) spindles in hermaphrodites, we observed lagging of both X chromosomes during anaphase I (**Fig. 2D**). These results reveal that anaphase I chromosome lagging during spermatocyte meiosis is likely caused by an inability to undergo synapsis rather than by a somatic effect of the male sex.

To further exclude that lagging is exclusive to the X chromosome, we analyzed hermaphrodites and males with the *zim-2*(tm574) mutation, which prevents the pairing of autosome IV (Phillips and Dernburg, 2006). We observed lagging chromosomes in all spindles in hermaphrodites (n = 10) and males (n = 5; **Fig. 2E and F**). Moreover, we created triploid males with spermatocytes containing five unpaired autosomes (Madl and Herman, 1979). In all meiosis I spindles of triploid worms (n = 32) we detected a massive fluorescent signal between segregating autosomes that correspond to the five unpaired autosomes (**Fig. 2G**). Collectively, these results show that lagging chromosomes during spermatocyte anaphase I are indeed a consequence of the lack of pairing and synapsis of any chromosome during prophase I and are not specific to sex chromosomes.

### Spermatocyte meiotic spindles display both anaphase A and B movement

We next investigated how spermatocyte meiotic spindles drive chromosome movement over time. We measured changes in the pole-to-pole (P-P), autosome-to-autosome (A-A) and pole-to-autosome (P-A) distances during both meiotic divisions using a strain with centrosomes labeled with γ-tubulin::GFP and chromosomes labeled with H2B::mCherry (**Fig. 3A-B**). In meiosis I, the pole-to-pole distance increased, which was evident in a sigmoidal curve where the initial and final spindle length are given by the two plateaus, and the segregation velocity was determined by the slope (**Fig. 3E**). The spindle length progressed from 4.1 ± 0.3 µm to 8.0 ± 0.6 µm (mean ± SD; n = 31) with an elongation speed of 1.29 ± 0.36 µm/min. Further, we found a simultaneous anaphase A-type movement in that the pole-to-autosome distance decreased by half, from 1.6 ± 0.3 µm to 0.8 ± 0.3 µm with a speed of −0.39 ± 0.27 µm/min (**Movie S2**; **Table 1**). Chromosome dynamics in meiosis II also exhibited anaphase A and B-type movements (**Fig. 3C-D and F, Movie S3; Table 1**).

**Figure 3.**
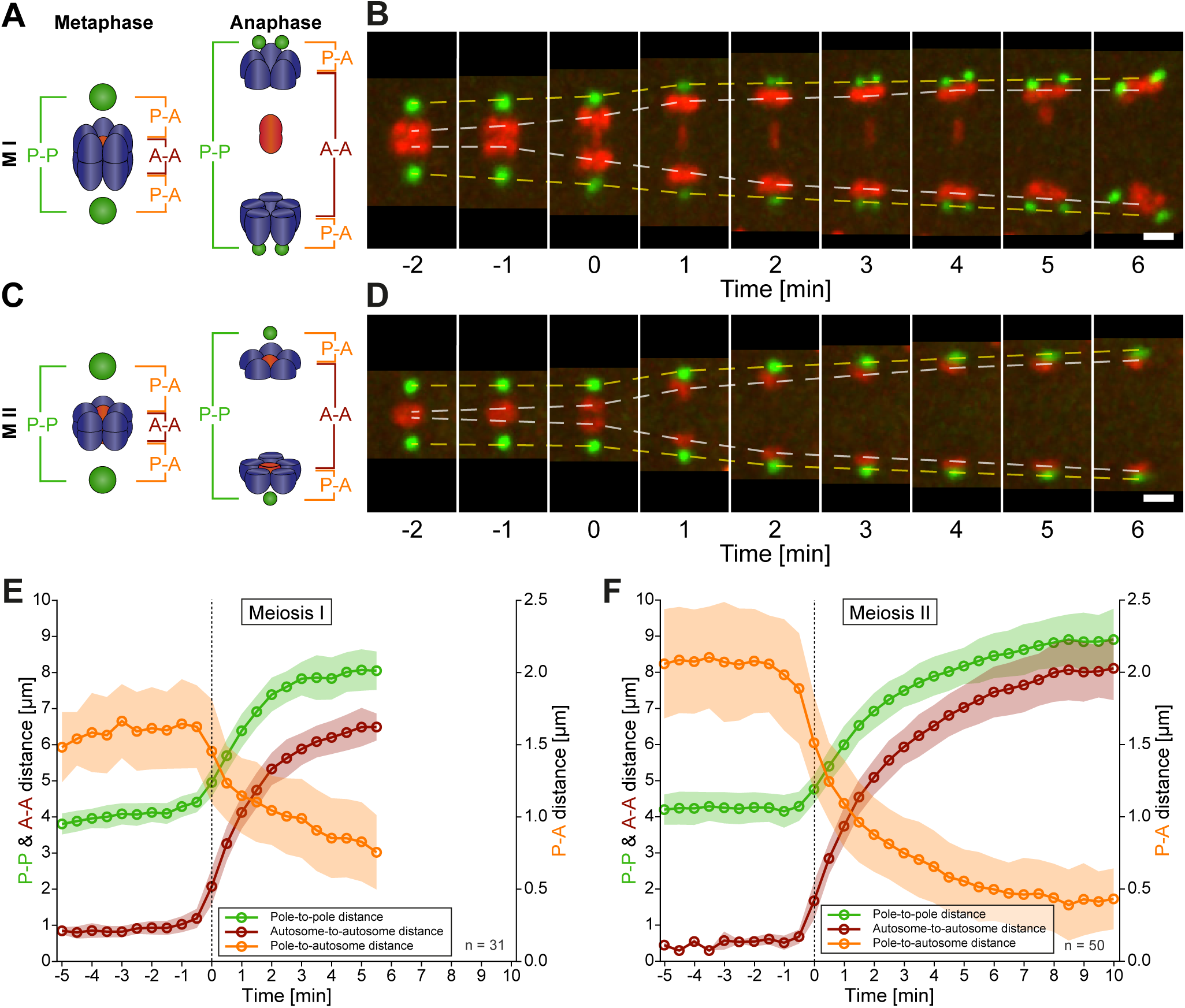
Spermatocyte meiotic spindles display both anaphase A and B movement. (**A**) Schematic representation of metaphase and anaphase during meiosis. Centrosomes are illustrated in green, autosomes in blue and the univalent X chromosome in red. The pole-to-pole (P-P, green), autosome-to-autosome (A-A, red), and both pole-to-autosome distances (P-A, orange) are indicated. (**B**) Time series of projections of a spindle in meiosis I obtained by confocal light microscopy. Centrosomes are labeled with γ-tubulin::GFP (green) and chromosomes with histone H2B::mCherry (red). The separation of the centrosomes (yellow dashed line) and autosomes (white dashed line) over time is indicated. Anaphase onset is time point zero (t=0). Scale bar, 2 µm. (**C**) Schematic representation of metaphase and anaphase during meiosis II as shown in (A). (**D**) Separation of centrosomes and autosomes in meiosis II as shown in (C). Scale bar, 2 µm. (**E**) Quantitative analysis of autosome and centrosome dynamics in meiosis I. Anaphase onset is time point zero (t=0). (**F**) Quantitative analysis of autosome and centrosome dynamics in meiosis II. The mean and standard deviation is given (circles and shaded areas) for (E) and (F).

Taken together, spermatocyte meiotic spindles in *C. elegans* exhibit both anaphase A and B-type movements. This is distinct from mitosis in the early *C. elegans* embryo, which utilizes only anaphase B mechanisms (Oegema et al., 2001), or the oocyte, which uses acentrosomal mechanisms during meiosis (Dumont et al., 2010; McNally et al., 2016; Muscat et al., 2015; Redemann et al., 2018). Similar to grasshopper spermatocytes (Ris, 1949), anaphase A and B movement occurs simultaneously, with anaphase A contributing approximately one fifth to the overall chromosome displacement in *C. elegans* spermatocyte meiosis.

### Spermatocyte meiotic centrosome and spindle dynamics are distinct from that in mitosis

To identify cell-type specific features of centrosomal segregation, we compared spindle pole dynamics in the two meiotic divisions in spermatocytes to those of the first mitotic division in one-cell embryos. During mitosis, centrosomes partially or completely disassemble after the single chromosome segregation event completes (Enos et al., 2018; Magescas et al., 2019). However, during the two rounds of spermatocyte meiosis, centrosomes must duplicate and migrate after sisters go to the same pole during meiosis I, then make new connections that allow sisters to segregate to opposite poles during meiosis II (Albertson and Thomson, 1993; Peters et al., 2010). Centrosome dynamics that accomplish this are not well understood. We tracked centrosome dynamics using γ-tubulin::GFP (Hannak et al., 2001; Strome et al., 2001) and found that centrosome size peaks at a volume of 3 µm^3^ at metaphase I (**Fig. S1A**) (Peters et al., 2010). After anaphase onset, the centrosome volume decreases rapidly to 1 µm^3^. At this point, the centrosomes flattened out before splitting into two distinct centrosomes (Schvarzstein et al., 2013), each at a volume of ∼1 µm^3^. These centrosomes then each migrate 90 degrees around the segregating autosomes to opposite poles, despite the absence of a nuclear envelope. Interestingly, the centrosomes remain connected to X chromosome-connected microtubules. After the X resolves and chromosomes align at metaphase II, the centrosome volume peaks at 1.3 µm^3^ then slowly decreases during anaphase II (**Fig. S1B**). Thus, the separation and migration of centrosomes during anaphase I appears to be coordinated with microtubules that must maintain connections not only to segregating autosomes, but also the lagging X chromosome.

We next compared chromosome and spindle dynamics of *C. elegans* spermatocytes to the one-cell embryo (**Fig. S2A, Table 1 and 2**) (Farhadifar et al., 2015). Despite the difference in cell and spindle size and structure, the final chromosome-to-chromosome distance of the spermatocyte meiotic spindle was 6.5 µm in meiosis I and 8.0 µm in meiosis II compared to the mitotic spindle, which is 8.5 µm (Nahaboo et al., 2015; Oegema et al., 2001). Spermatocyte meiosis I is slower, completing in 4-5 minutes compared to mitotic division, which is accomplished over 1.5 min (**Table 2**). Thus, chromosomes during the first mitotic division in *C. elegans* segregate three times faster (6.5-7.5 µm/min) than in spermatocyte meiosis I (2.07 µm/min, n = 31) (Nahaboo et al., 2015; Oegema et al., 2001). During mitotic anaphase, a mid-spindle structure, the central spindle, forms between the segregating chromosomes (Yu et al., 2019). In contrast, during spermatocyte meiosis, the lagging X chromosome precludes detection of a similar inter-chromosomal array of microtubules (**Fig. S3A**). To visualize non-X-connected mid-spindle microtubule structures, we examined spermatocytes in *tra-2*(e1094) (XX) animals, where paired X chromosomes do not lag. We detected only a weak fluorescence signal in between the segregating chromosomes (**Fig. S3B**), not indicative of the prominent central spindle structures observed in mitosis (**Fig. S2A**). Overall, we find that the spermatocyte meiotic spindle, though smaller and slower than mitotic spindles, can move chromosomes a similar distance.

### Meiotic spindle dynamics are faster in spermatocytes than oocytes

We found spindle dynamics differ dramatically between spermatocytes and oocytes. Meiotic spindles in *C. elegans* are similarly sized in spermatocytes and oocytes, even though oocyte meiotic spindles are acentrosomal (Albertson and Thomson, 1993; Dumont et al., 2010; McNally et al., 2016; Muscat et al., 2015; Redemann et al., 2018). At metaphase the female spindles are positioned near the embryonic cortex (Fabritius et al., 2011) and are 6-7 µm in length (**Fig. S2B, Table 2**). At anaphase onset, a shortening of pole-to-chromosome distance is observed before a central array of microtubules forms in between segregating chromosomes to push them apart to a final distance of 4.5-6 µm (Dumont et al., 2010; Laband et al., 2017; McNally et al., 2016; Muscat et al., 2015; Redemann et al., 2018; Yu et al., 2019). Both female meiotic divisions segregated at comparable rates of 0.6-0.8 µm/min (McNally et al., 2016), which is significantly slower than spermatocyte meiotic chromosome movement (1.29 µm/min). Thus, though spermatocyte and oocyte spindles are comparably sized, centrosomal spermatocyte spindles segregate chromosomes twice as fast over a longer distance.

### Kinetochores are not disassembled between spermatocyte meiotic divisions

Kinetochores establish critical connections between chromosomes and microtubules that are required for chromosome movement. Previous studies in mitosis and oocyte meiosis found the inner kinetochore proteins CENP-A^HCP-3^ (Maddox et al., 2012) and CENP-C^HCP-4^ (Moore and Roth, 2001) form a platform on chromosomes that connects to the outer kinetochore, composed of the MIS-12 complex (Kline et al., 2006; Petrovic et al., 2016; Petrovic et al., 2010), KNL-1 (Desai et al., 2003; Ghongane et al., 2014), and the NDC-80 complex (Cheerambathur et al., 2017; Kudalkar et al., 2015; Wilson-Kubalek et al., 2016), which interacts with microtubules (Cheeseman et al., 2006; Ciferri et al., 2008; Wei et al., 2007) (**Fig. 4A**). Kinetochore components can localize on, surround, or cup the ends of rounded oocyte meiotic chromosomes (**Fig. 4B**) (Dumont et al., 2010; McNally et al., 2016; Muscat et al., 2015; Shakes et al., 2009). To identify sperm-specific kinetochore dynamics, we analyzed kinetochore protein localization in fixed spermatocytes compared to oocytes (**Fig. 4C**).

**Figure 4.**
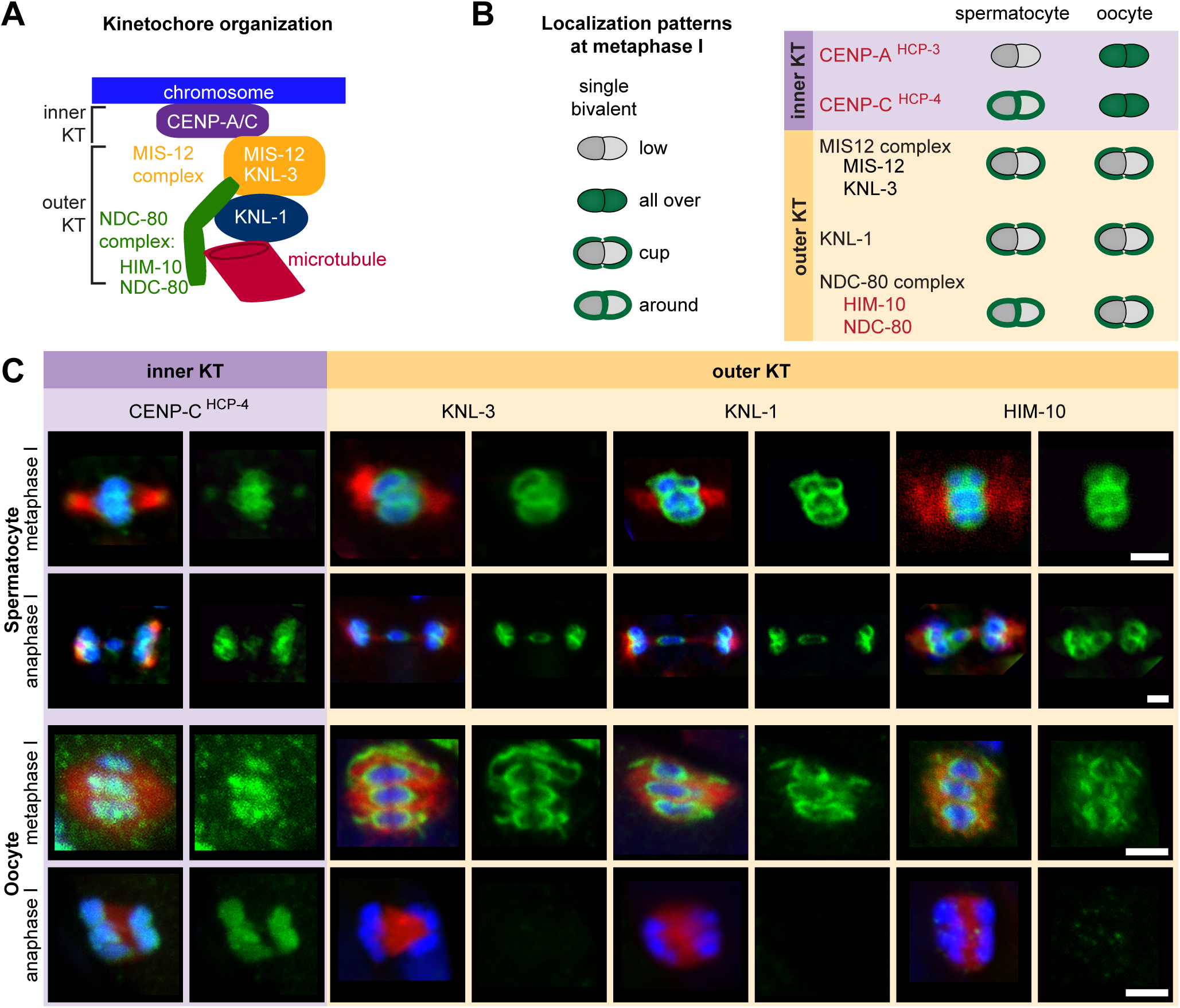
Kinetochores do not disassemble between spermatocyte meiotic divisions. (**A**) Schematic representation of kinetochore (KT) organization in *C. elegans*. (**B**) Localization patterns of kinetochore proteins on single bivalents (schematic illustration, left panel; summary of immunostainings as shown in (C), right panel). Kinetochore proteins with a similar pattern of localization in sperm and oocytes are listed in black, proteins with a gamete-specific pattern in red. (**C**) Visualization of kinetochore proteins in sperm (upper two panels) and oocyte meiosis I (lower two panels). Metaphase (upper row in each panel) and anaphase (lower row in each panel) of the first division is given. Males were fixed and stained with antibodies against kinetochore proteins (green), microtubules (red), and DAPI (blue) and imaged by confocal light microscopy. For each kinetochore protein, the left panels show whole spindles with microtubules, chromosomes and kinetochores; right panels show the localization patterns of the kinetochore protein only. Scale bars, 2 µm.

During sperm metaphase I and anaphase I, the inner kinetochore protein CENP-C^HCP-4^ is more concentrated surrounding chromosomes (**Fig. 4B-C**), and CENP-A^HCP-3^ is largely absent (Shakes et al., 2009). In contrast, on oocyte chromosomes CENP-A^HCP-3^ and CENP-C^HCP-4^ are both present and uniformly spread on chromosomes (Monen et al., 2005; Shakes et al., 2009). The outer kinetochore proteins, KNL-1 and KNL-3, localize around sperm chromosomes in metaphase I (**Fig. 4B-C)**, but cup the poleward ends of chromosomes during oocyte meiosis (Dumont et al., 2010). Further, HIM-10, a member of the NDC-80 complex, surrounds chromosomes at metaphase I and anaphase I, unlike in oocyte meiosis, where it cups chromosome ends. Strikingly, during oocyte anaphase I and II, when outer kinetochore protein levels are greatly reduced, they are retained throughout spermatocyte divisions (**Fig. 4B-C**). Thus, both inner and outer kinetochore proteins show sperm-specific localization and dynamics, particularly the retention of outer kinetochore components between meiotic divisions.

### Microtubules attached to the X chromosome exert a pulling force

The retention of kinetochore components on poleward sides of chromosomes during anaphase suggests that kinetochore-attached microtubules could contribute to pulling chromosomes. To investigate this, we hypothesized that we could detect tension that would alter the shape of the lagging X as evidence for pulling forces acting on chromosomes during anaphase separation (**Fig. 5A**). Using a strain expressing histone H2B::mCherry, we observed the X chromosome was stretched along the spindle axis in early anaphase. As X resolves to one side at late anaphase, it rounds up (**Fig. 5A**, upper panel). To quantify changes in chromosome geometry, we calculated a shape coefficient (the ratio of length over diameter) of the X chromosome over time (**Fig. 5A**, lower panel). With this measure, stretched chromosomes have a shape coefficient greater than 1. Indeed, the X chromosome is significantly stretched early in anaphase I with a shape coefficient of 1.4 that decreases to 1.0 as it rounds up in late anaphase I (**Fig. 5B**). This suggests that the X chromosome is under tension during anaphase I, which is released as the lagging chromosome resolves to one side.

**Figure 5.**
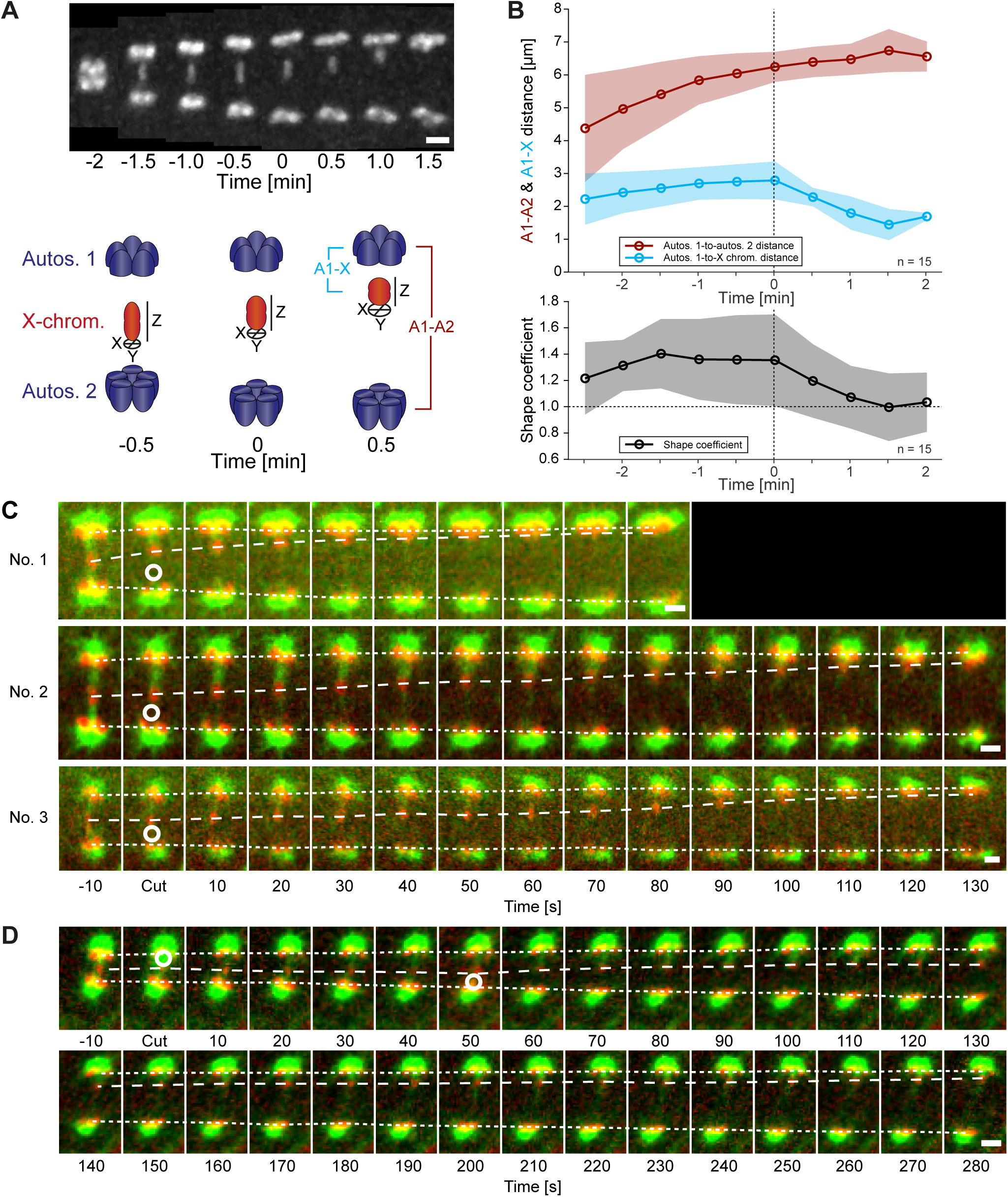
Microtubules attached to the X chromosome exert a pulling force. (**A**) Maximum intensity projection images with chromosomes labeled with histone H2B::mCherry (upper panel, shown in white). Time is relative to the onset of segregation of the X chromosome (t=0). Scale bar, 2 µm. Illustration of X chromosome shape quantitation (lower panel). The length (Z) of the X chromosome (red) is divided by its width (mean of X + Y). Autosomes are shown in blue. Time is relative to the onset of segregation of the X chromosome (t=0). (**B**) Plot showing the shape of chromosomes in male spindles (n=15). The upper panel shows the autosome 1-to-autosome 2 (A-A, blue) and the autosome 1-to-X chromosome distances (A-X, red) over time, the lower panel the shape coefficient (black). Solid lines show the mean, shaded areas indicate the standard deviation. Time zero (t=0) is the onset of X chromosome movement. (**C**) Microsurgery of microtubules associated with the X chromosome in anaphase I. Microtubules are labeled with β-tubulin::GFP (green) and chromosomes with histone H2B::mCherry (red). Time is given relative to the time point of the applied laser cut (t=0). The position of the cut is indicated (white circle). The position of the autosomes (outer dashed lines) and the X chromosome (inner dashed line) is indicated. The three panels show examples with a fast (top), intermediate (middle) and slow response (bottom) of X chromosome segregation to the applied cut. Scale bars, 2 µm. (**D**) Example of a double cut experiment. The panels show a single experiment over about 300 s. The two cuts are indicated (white circles). Scale bar, 2 µm.

To further assess pulling forces during anaphase, we used laser microsurgery on X chromosome-attached microtubules. We reasoned a cut on one side of the lagging X would release tension and induce segregation to the opposite side. Using a β-tubulin::GFP and histone H2B::mCherry expressing strain, we applied laser point-ablations to the microtubule bundle on one side of the lagging X. We observed an immediate and continuous movement of the X chromosome towards the unablated side (**Fig. 5C**, **Movie S4**). The velocity of X chromosome movement after the cut was variable, similar to unperturbed spindles (**Fig. 5C**, see also cut no. 3). We also tested cutting the microtubule bundles sequentially on each side of the X chromosome. After initiation of movement by the first cut, the second cut on the opposite side led to a rapid change in the direction of segregation (**Fig. 5D**, **Movie S5**), indicating that the direction of X movement can be reversed and that microtubule connections are highly dynamic during anaphase I.

Taken together, we conclude that kinetochore microtubules exert a pulling force on chromosomes during anaphase. This tension is most obvious on the microtubules connected to X chromosome, which eventually causes attachments to stochastically break as poles separate, allowing the X to resolve to the side with more attached microtubules.

### Spermatocyte meiotic spindle structure at metaphase is distinct from that in oocyte meiosis

To investigate how microtubules connect to chromosomes during spermatocyte meiosis, we used large-scale electron tomography (Redemann et al., 2017; Redemann et al., 2018) to visualize the ultrastructure of whole meiotic spindles with single-microtubule resolution. We generated 3D models of one spindle at metaphase (**Fig. 6A**), one spindle at anaphase onset (**Fig. 6B**) and four spindles with increasing pole-to-pole length at anaphase (**Fig. 6C-F**, referred to as anaphase no. 1-4). For each reconstruction we segmented the autosomes (a), the X chromosome (x), and the centrioles (**Fig. 6**, left panels). Using criteria applied to previously described mitotic holocentric kinetochores, microtubules within a ribosome-free zone of 150 nm around chromosomes were considered kinetochore microtubules (O’Toole et al., 2003; Redemann et al., 2017). All other microtubules were annotated as spindle microtubules (**Fig. 6**, mid left panels; mid right panels showing kinetochore microtubules only; **Table 3**). This distance is in accord with the interaction distance measured by super-resolution light microscopy (**Fig. S4**).

**Figure 6.**
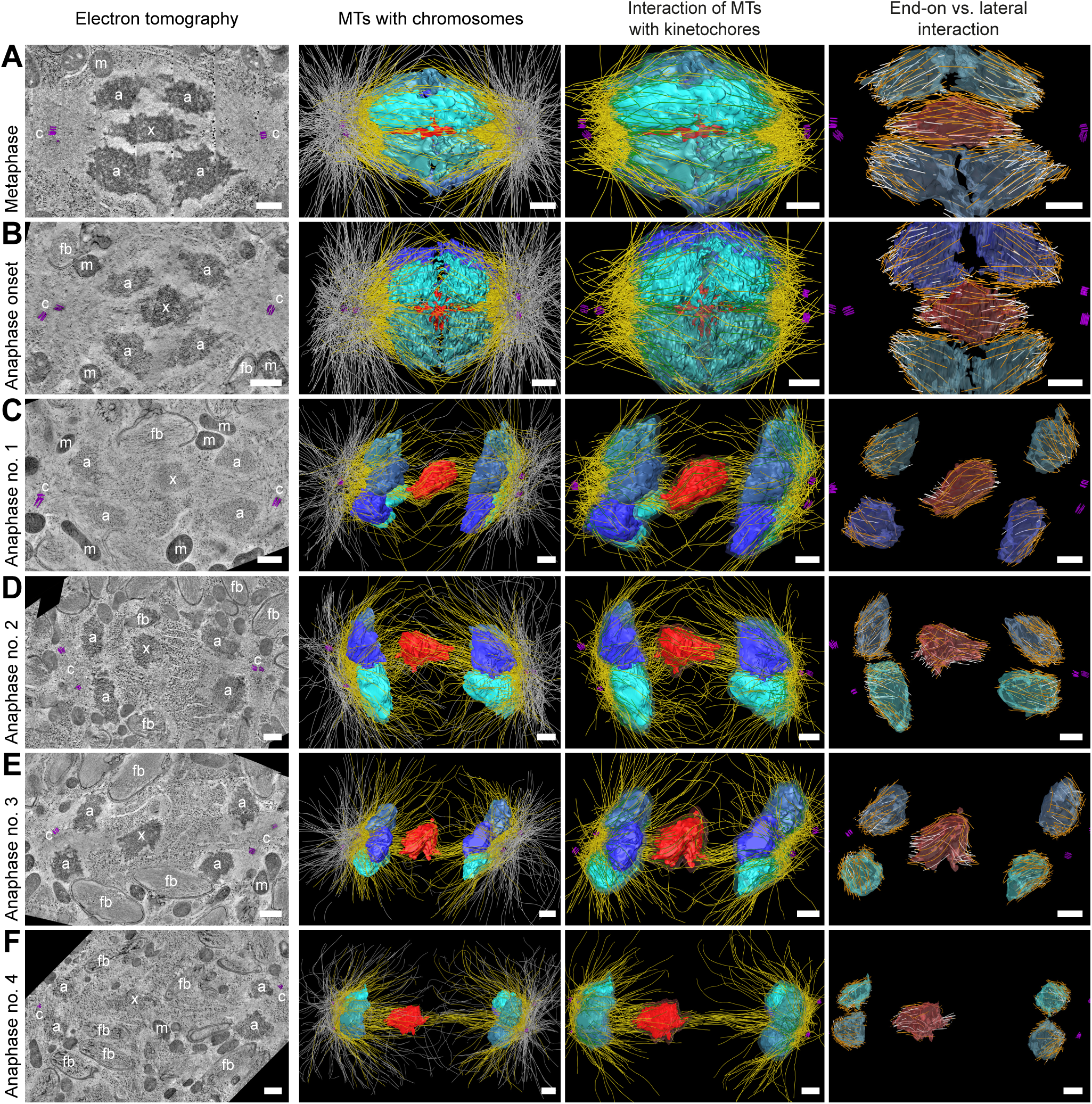
Three-dimensional ultrastructure of spindles in spermatocyte meiosis I. Full tomographic reconstruction of spermatocyte spindles. Scale bars, 500 nm. (**A**) Metaphase spindle. (**B**) Spindle at anaphase onset. (**C**) Spindle at mid anaphase with a pole-to-pole distance of 2.98 µm. (**D**) Mid anaphase spindle with a pole-to-pole distance of 3.35 µm. (**E**) Mid anaphase spindle with a pole-to-pole distance of 3.37 µm. (**F**) Spindle at late anaphase with a pole-to-pole distance of 5.45 µm and the X chromosome with initial segregation to one of the daughter cells. Left panels: tomographic slice showing the centrosomes (colored purple, c), the autosomes (a), and the unpaired X chromosome (x) aligned along the spindle axis. Mitochondria (m) and fibrous body-membranous organelles (fb) are also indicated. Mid left panels: corresponding three-dimensional model illustrating the organization of the full spindle. Autosomes are shown in different shades of either blue or cyan, the X chromosome in red, centriolar microtubules in purple, microtubules within 150 nm to the chromosome surfaces in yellow, and all other microtubules in gray. Mid right panels: interaction of microtubules with the kinetochores. Kinetochores are shown as semi-transparent regions around each chromosome. The part of each microtubule entering the kinetochore region around the holocentric chromosomes is shown in green. Right panels: visualization of end-on (white) *versus* lateral (orange) interactions of microtubules with chromosomes. Only the parts of microtubules inside of the kinetochore region are shown.

Though similarly-sized, round sperm and oocyte chromosomes both adopt a rosette arrangement surrounding the X chromosome, the meiotic spindle in spermatocytes is distinct from that of oocytes. During spermatocyte meiotic metaphase I, of the 2406 total number of microtubules that compose the spindle, 912 (38%) were kinetochore microtubules and 393 (16%) made end-on attachments to chromosomes. This is in contrast to oocyte metaphase meiotic spindles, which are composed of ∼1/3 more microtubules (3662-3812 microtubules) (Redemann et al., 2018). Of these, 1038-1402 (∼32.5%) were kinetochore microtubules and only 131-165 (∼4%) of which made end-on attachments. This indicates that round chromosomes can both be aligned at metaphase in sperm centrosomal and oocyte acentrosomal spindles but using significantly different ratios of lateral and end-on attachments.

### Continuous and lengthening microtubules connect the X chromosome to centrosomes during anaphase I

One phenomenon of spermatocyte meiosis is that microtubules, which connect the chromatids of the lagging X to opposite spindle poles, lengthen during anaphase I. This is unusual because in most centrosomal cell-types, microtubules either shorten (anaphase A) or stay the same length as poles are pulled apart (anaphase B). Further, in *C. elegans* mitosis, continuous microtubules do not directly connect between centrosomes and chromosomes, but instead anchor into the spindle network (Redemann et al., 2017). We thus used electron tomography to determine the continuity of individual X-connected kinetochore microtubules during anaphase I. We found that microtubules directly connect the X chromosome to each centrosome throughout anaphase I (see also **Fig. 6A-F**, mid left panels; **Fig. S5**). Further, microtubules with both end-on and lateral interactions to X remained continuous even as they increased in length until the X chromosome resolved to one side (**Fig. 6A-F**, right panels; **Fig. S5A-C**, right panels; **Fig. 7A-B; Fig. S6**). Thus continuous microtubules connect the X chromosome to poles even as poles elongate during anaphase I.

**Figure 7.**
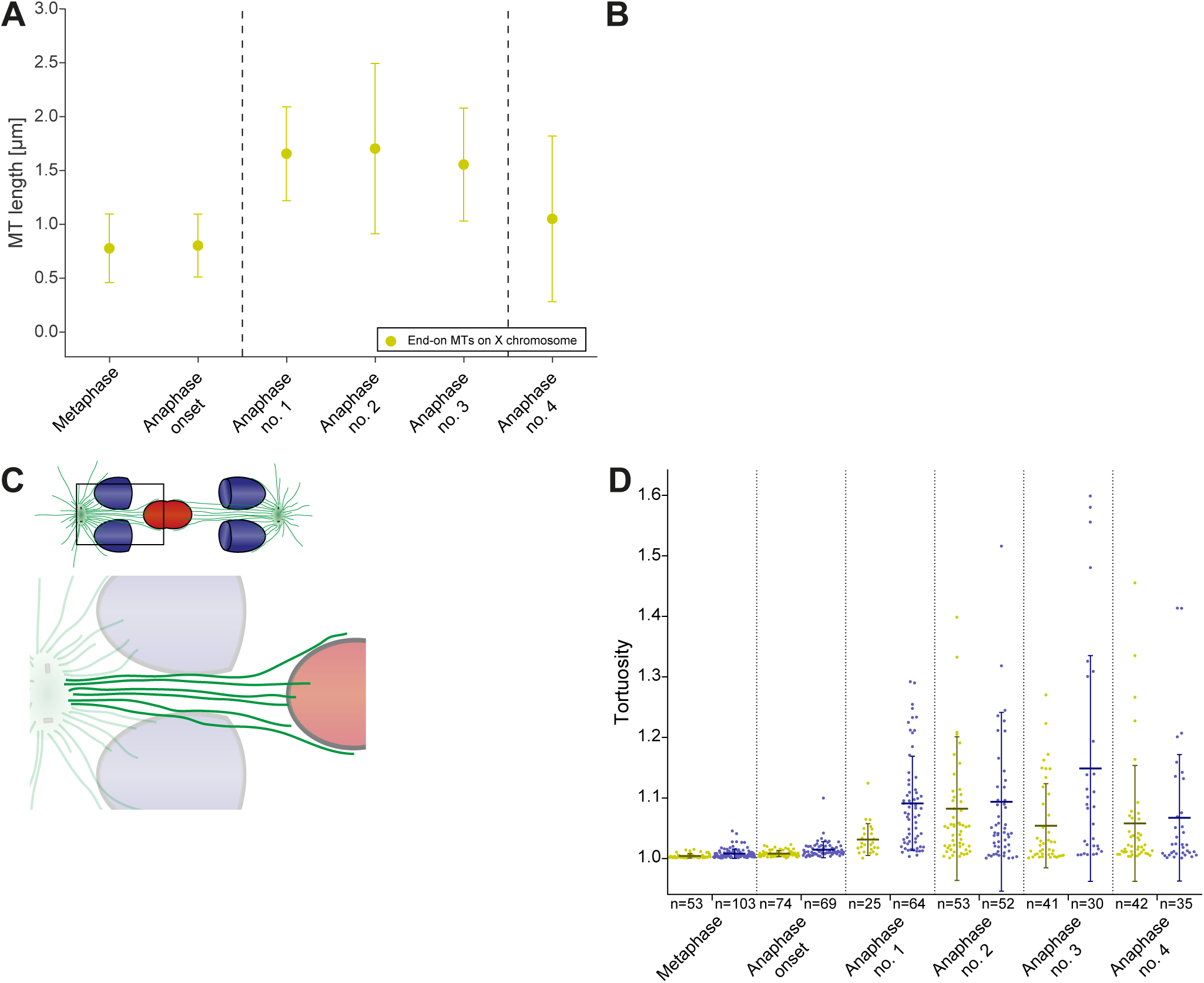
X chromosome-attached microtubules are continuous and lengthen during anaphase I. (**A**) Length distribution of end-on X chromosome-attached microtubules at different stages of meiosis I (corresponding to data sets as shown in Fig. 6). Dots show the mean, error bars indicate the standard deviation. (**B**) Length distribution of laterally X chromosome-attached microtubules at different stages of meiosis I. (**C**) Schematic drawing illustrating X chromosome-attached kinetochore microtubules. Both end-on (yellow) and laterally attached microtubules (purple) are shown (left panels). Curvature of individual microtubules is measured as illustrated (right panel) using tortuosity, which is spline length (red dotted lines) divided by end-to-end length (black dotted lines). (**D**) Plot showing the tortuosity of end-on and laterally attached kinetochore microtubules. The meiotic stages correspond to the data sets as shown in Figure 6. The mean, standard deviation and individual measurements are given for each data set.

We also observed that X-connected microtubules were curved during late anaphase I. To investigate this, we determined the curvature of X-connected end-on and laterally attached kinetochore microtubules by measuring the tortuosity of individual microtubules. For this, we calculated the ratio of the spline length over the end-to-end length (**Fig. 7C**). At metaphase and anaphase onset, kinetochore microtubules had a tortuosity ratio of one, indicating that the microtubules were straight. In contrast, X-connected microtubules exhibited higher tortuosity at anaphase, indicating a higher curvature (**Fig. 7D**). This suggests that other cellular forces, besides those generated by pulling forces, may also be acting on microtubules connected to the lagging X during anaphase I.

### Segregation of the X chromosome correlates with an asymmetry in the number of attached microtubules

To further characterize the nature of X chromosome lagging and resolution, we examined tomographic data to determine the total microtubule length in confined volumes on each side of the X (**Fig. 8A-B**). We determined the ratio of the total microtubule length for each tomographic data set, which were then plotted according to autosome-to-autosome distance (**Fig. S7A-B**). We further determined the number of kinetochore microtubules associated with the opposite hemispheres of the X chromosome and calculated the ratio of these two values (**Fig. 8C-D**). The ratio of total microtubule length and the ratio of microtubule number is about one in metaphase and early anaphase, suggesting microtubules are present equally on both sides. As anaphase progresses, this ratio deviates from one, indicating less microtubules on one of the two sides, presumably enabling the X chromosome to resolve to the opposing side.

**Figure 8.**
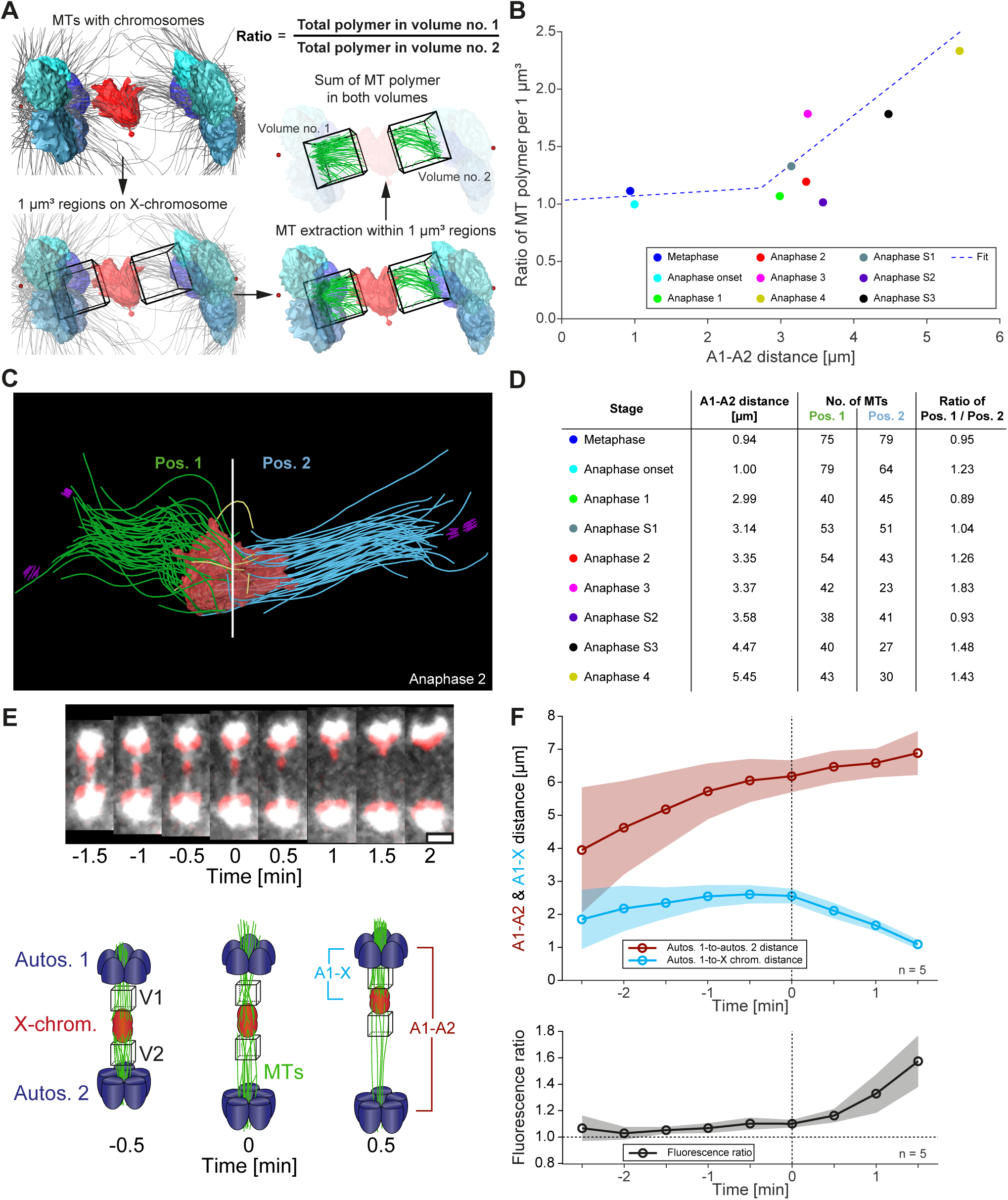
Resolution of the X chromosome to one side correlates with an asymmetry of microtubules. (**A**) Left: EM model of an anaphase I spindle and definition of two identical volumes at opposite positions to the X chromosome. Right: Extraction of microtubules and measurement of the total polymer length within the selected volumes. The total length of microtubules was measured within volumes of 1 µm^3^. (**B**) Graph showing the ratio of both volumetric polymer length measurements plotted against the autosome-autosome distance for each individual data set corresponding to the stages shown in Fig. 6 and Fig. S5. A trend line was fitted to illustrate the increase in the asymmetry. (**C**) Deconstructed 3D model (anaphase 2 data set) illustrating the total number of microtubules attached on each side of the X chromosome (red), named pos. 1 (microtubules shown in green) and pos. 2 (microtubules shown in blue). Centrioles are shown in purple, undefined microtubules in yellow. (**D**) Table showing the autosome 1-to-autosome 2 distance (A1-A2), the total number of microtubules for both positions and the calculated ratio for each data set. (**E**) Upper panel: Maximum intensity projection images from live imaging show microtubules attachments to the segregating X chromosome. Microtubules are labeled with β-tubulin::GFP (white) and chromosomes with histone H2B::mCherry (red). Time is given relative to the onset of X chromosome segregation (t=0). Scale bar, 2 µm. Lower panel: Illustration of the measurement of fluorescence intensity in two volumes (V1, V2) of 1 µm^3^ each at opposite sides of the X chromosome (red), autosomes (blue), microtubules (green). (**F**) Ratio of fluorescence intensities (V1/V2) as illustrated in (A). Upper panel: the autosomes 1-to-autosomes 2 distance (A-A, red) and the autosomes 1-to-X chromosome distance (A-X, blue) over time. Solid lines show the mean, shaded areas indicate the standard deviation. Time zero (t=0) corresponds to the onset of X chromosome movement. Lower panel: the ratio of fluorescence intensities is given for corresponding time points (black, time is relative to the onset of segregation of the X chromosome, t=0; n=5).

To confirm that differences in microtubule content we observed by EM correlate with X resolution, we tracked microtubules (β-tubulin::GFP) relative to chromosomes (histone H2B::mCherry) by live-cell imaging (**Fig. 8E**, upper panel). The sum of GFP fluorescence was measured in a similar volume on each side of the X chromosome over time (**Fig. 8E**, lower panel). The ratio of the two volumes enabled us to indirectly quantify microtubule content differences on each side of X. During early anaphase I, the ratio of the volumes was almost one, indicating a similar amount of microtubules connected to each side. As the X chromosome segregated to one side over the other, we detected increased intensity on the side the X moved closer toward (**Fig. 8F**). This indicates an asymmetry in attached microtubules correlates with X chromosome resolution.

### Autosomal attached kinetochore microtubules do not shorten during anaphase A

Our live imaging and EM data both show an anaphase A autosome-to-centrosome distance decrease during anaphase I (**Table 3**). Because a well-described mechanism for anaphase A (chromosome-to-pole shortening) is microtubule shortening (Asbury, 2017), we analyzed individual kinetochore microtubule lengths in our 3D EM reconstructions during anaphase. At metaphase, end-on kinetochore microtubules attached to autosomes were 0.62 ± 0.33 µm in length (n = 360) (**Fig. 9A**), while laterally attached microtubules were much longer, at 1.15 ± 0.59 µm (n = 480) (**Fig. 9B**). Unexpectedly, we observed that end-on kinetochore microtubules did not shorten as anaphase I progressed, remaining at 0.63 ± 0.42 µm (n = 687). Thus, unlike in other systems, the anaphase A we observe in spermatocyte meiosis is not due to shortening of end-on attached kinetochore microtubules (Asbury, 2017).

**Figure 9.**
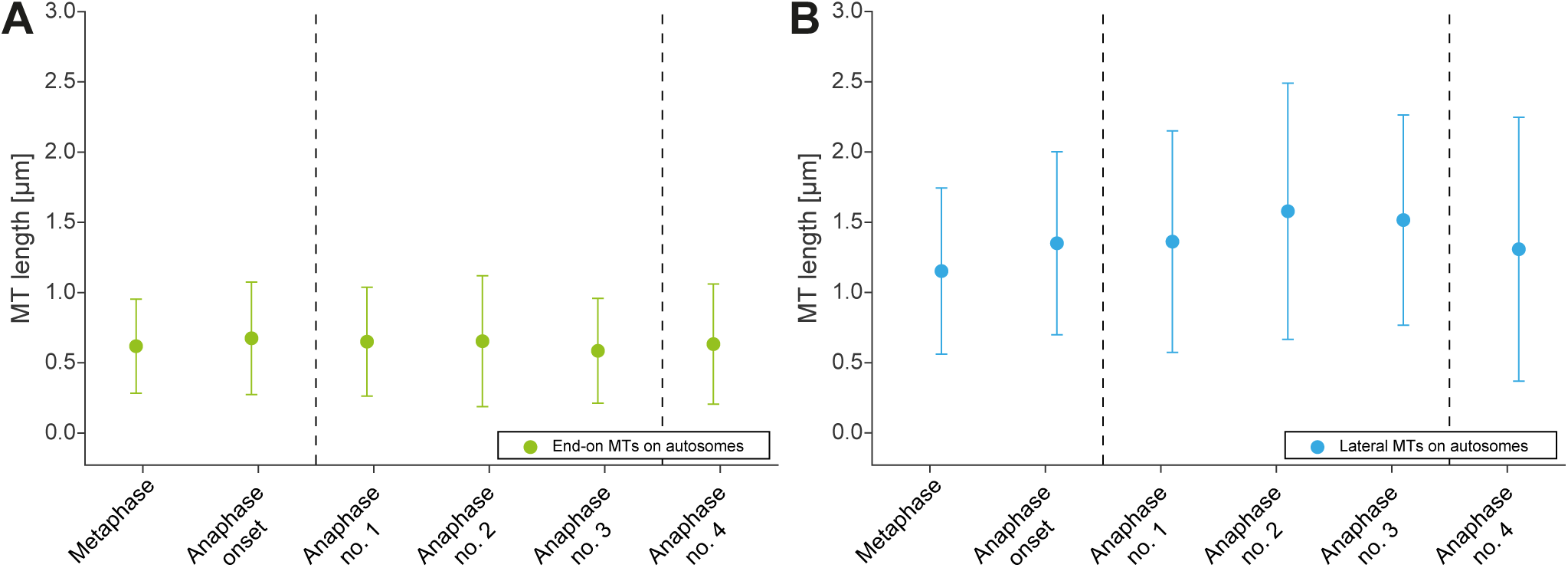
Autosome-attached kinetochore microtubules do not shorten during anaphase. (**A**) Length distribution of end-on autosome-attached microtubules at different stages of meiosis I (corresponding to data sets as shown in Fig. 6). Dots show the mean, error bars indicate the standard deviation. (**B**) Length distribution of laterally autosome-attached microtubules at different stages of meiosis I.

### Tension release across the spindle may contribute to autosomal anaphase A

To account for anaphase A in spermatocyte meiosis, we hypothesized that features of spindle geometry and shape could contribute to the decrease in chromosome-to-centrosome distance. We thus analyzed changes in the shape of chromosomes, centrosomes, and the interaction angles of kinetochore microtubules with the autosomes over the course of anaphase that could contribute to this decrease.

First, to examine the contribution of chromosome stretch, we measured individual autosome expansion along the spindle axis by plotting the cross-sectional areas over the chromosome distance. This generates a stretch value we call the Full Width at Half-Maximal or FWHM (**Fig. 10A**; see experimental procedures). Autosomes are stretched most at metaphase (FWHM: 0.73 ± 0.13 µm; n = 10) (**Fig. 10B**). As chromosomes separate farther apart, autosomes round up to a FWHM of 0.56 ± 0.06 µm, about 23% less compared to metaphase, thereby moving the chromosome centers closer to the poles. Thus, the stretching of chromosomes induced by metaphase alignment that is released during anaphase progression accounts for a portion of anaphase A pole-chromosome shortening.

**Figure 10.**
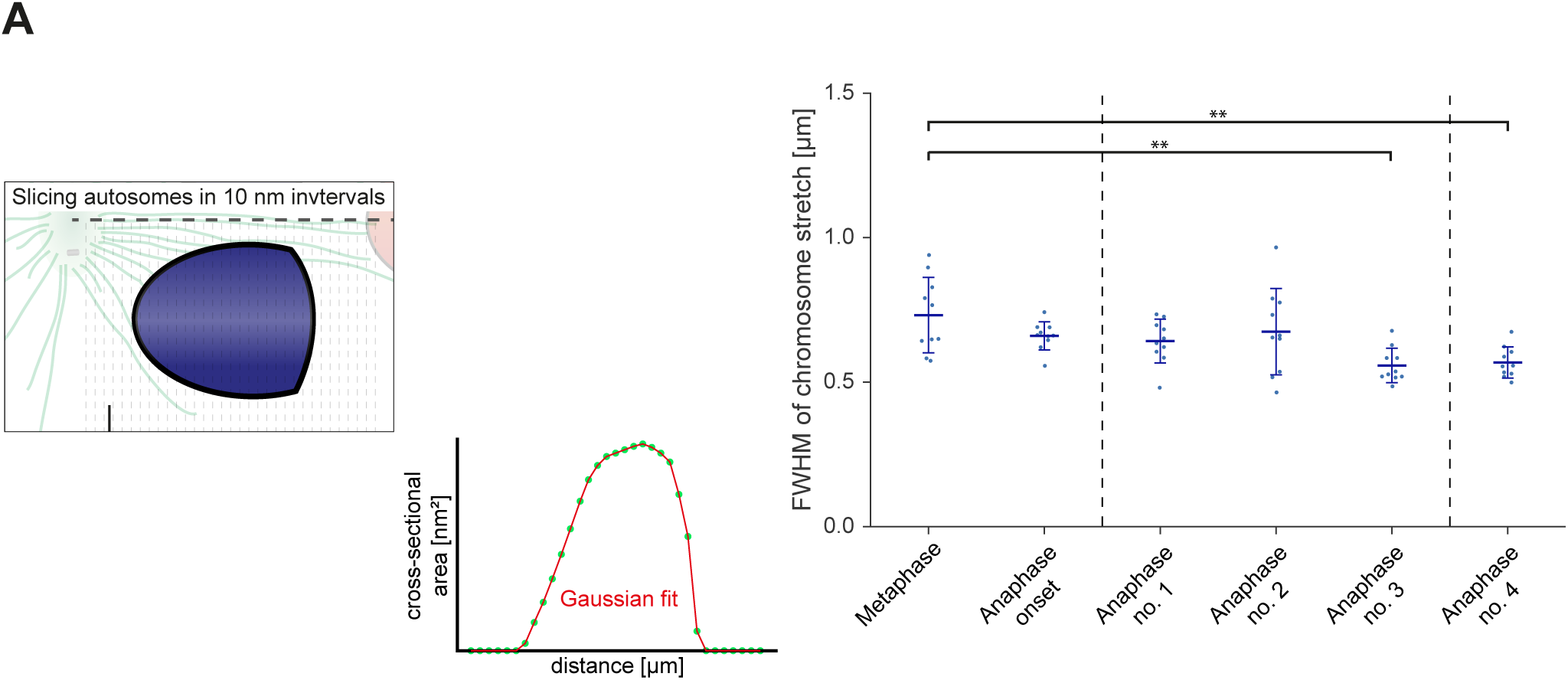
Changes in the geometry of spindles account for anaphase A pole-chromosome shortening. (**A**) Analysis of autosome stretch. Schematic representations of anaphase I showing the determination of stretch along the pole-to-pole spindle axis for an individual autosome (left panels). Stretch is indicated by the full width half maximum (FWHM), which was quantified as shown in the right panels. (**B**) Plot showing the FWHM of chromosome stretch for each meiotic stage analyzed. The mean, standard deviation and single measurements (n = 10) for each meiotic stage (corresponding to the ones as shown in Fig. 6) are given. One-way analysis of variance (ANOVA) of the metaphase dataset against anaphase datasets no. 3 and 4 are shown. Level of significance: ** represents p <= 0.01. (**C**) Analysis of the distance of individual kinetochore microtubule plus ends to the closest centriole (left panels: schematic drawings). For each kinetochore microtubule (green line), the direct distance (yellow line) from the putative plus end to the respective centriole was measured (right panels; plus ends of kinetochore microtubules are circles). (**D**) Plot showing the distance of kinetochore microtubule plus-ends to centrioles at different meiotic stages as described in (B). The mean, standard deviation and single measurements for each meiotic stage are given. One-way ANOVA of the metaphase dataset against anaphase datasets no. 3 and 4 are shown. Levels of significance: * is p <= 0.05; and *** is p <= 0.001. Additional ANOVA results are in Fig. S8H. (**E**) Analysis of the attachment angle of kinetochore microtubules associated end-on to autosomes. The schematic illustrates the defined main axis for the measurements (left panel: dashed line from the center of each autosome to the center of the centrosome,). Right panel: the angle (□) between each line connecting the kinetochore microtubule plus-end and the autosome center (green lines) and the main axis (dashed line) was measured for each kinetochore microtubule. (**F**) Plot showing the angle measurements from (E). The mean, standard deviation and single measurements for each meiotic stage are given. One-way ANOVA of the metaphase dataset against anaphase datasets no. 3 and 4 are shown. Levels of significance: *** is p <= 0.001. Additional ANOVA results are in Fig. S8I.

Second, we considered the distance between chromosomes and the centriole as centrosomes change shape as it splits (**Fig. S2**). The shift from a spherical to a stretched organelle could account for spindle poles moving closer to chromosomes. We measured the distance of the plus end of the kinetochore microtubules to the closest centriole and found it significantly shortened from metaphase (0.99 ± 0.27 µm) to anaphase (0.79 ± 0.21 µm) (**Fig. 10C**), resulting in autosomes being 20% closer to centrioles (**Fig. 10D**).

Third, we hypothesized tension release would also alter the angle of attachment of end-on kinetochore microtubules with autosomes, bringing chromosomes closer to spindle poles. We thus determined the attachment angle between each kinetochore microtubule plus-end at the chromosome surface and each centrosome-chromosome axis (**Fig. 10E**). The angle at metaphase was 36.8° ± 31.0°. Concomitant with a rounding up of the autosomes, the attachment angle increased to 58.8° ± 33.8°, bringing chromosomes closer to poles during anaphase (**Fig. 10F**). Simple trigonometric calculations with a constant microtubule length of 0.63 µm found this increase contributes to 0.17 µm shortening in chromosome to pole distance.

In sum, we developed analyses to identify three different factors that can contribute to pole-chromosomes shortening during anaphase: 1) the loss of tension at the chromosomes after anaphase onset, which shortens the chromosome by about 0.34 µm; 2) changes in centrosome size and shape that contributes about 0.2 µm; and 3) the opening of the attachment angle that is about 0.17 µm. All these factors comprise about ∼70% of the total ∼1 µm shortening of the chromosome-to-pole distance observed in spermatocyte meiosis (**Fig. S8A-I**), though this may be underestimated due to limitations in the tomographic reconstruction from serial semi-thick sections. Overall, our ultrastructure analysis revealed previously unknown, alternative mechanisms that contribute to anaphase-A movement.

## Discussion

Prior to this work, few studies addressed the spindle architecture that segregates chromosomes during male meiosis, a fundamental factor in sperm production (LaFountain et al., 2011; LaFountain et al., 2012; Nicklas and Kubai, 1985; Nicklas et al., 2001; Zhang and Nicklas, 1995). Here the combination of live imaging and 3D ultrastructure in *C. elegans* enabled us to define specific features of chromosome dynamics and spindle organization that are regulated in sex-specific ways to produce different cell types.

### A distinct form of anaphase A without shortening of kinetochore microtubules

Surprisingly, the single-microtubule resolution of electron tomography revealed the length of autosome-attached kinetochore microtubules is constant during sperm anaphase. This is in contrast to the kinetochore microtubule shortening typically associated with anaphase A observed by light microscopy in many systems (Asbury, 2017). We developed methods to analyze ultrastructural data to identify three contributors to the anaphase A in *C. elegans* spermatocyte meiosis (**Fig. 11A**). First, stretched autosomes at metaphase relax from tension released by separase-mediated cleavage of cohesins during anaphase, resulting in autosome shape change (Severson and Meyer, 2014). Second, live imaging revealed spindle poles decrease in size and change shape as centrioles split. Analysis of our ultrastructural data revealed that this results in a shorter distance between microtubule ends and the centrioles. Third, microtubule ends on round chromosomes shift from a central to more peripheral position during anaphase, resulting in a decrease in the microtubule-to-pole distance. This might be induced by minus-end associated motor proteins such as dynein (Schmidt et al., 2005; Schmidt et al., 2017). All three factors change the relative position of centrosomes, autosomes and kinetochore microtubules to one another. Such changes in the relative positioning without kinetochore microtubule shortening represent a novel type of centrosomal anaphase A movement that can now be considered when analyzing chromosome movement in different system and interrogated by the methods we describe.

**Figure 11.**
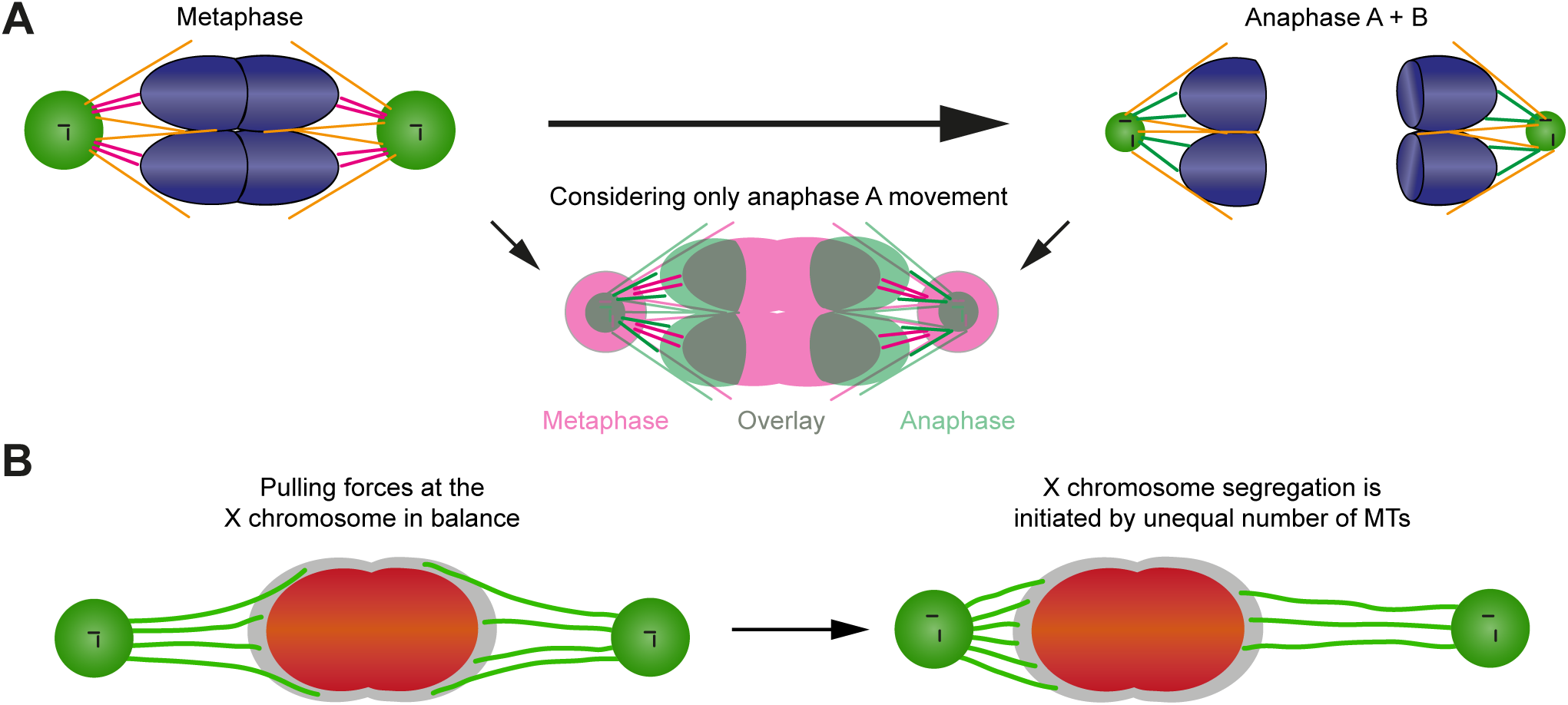
Proposed models of chromosome movements in meiosis I. (**A**) Model illustrating the anaphase movement of autosomes in sperm meiosis in the absence of kinetochore microtubule shortening. Left upper panel is metaphase; right upper panel is anaphase; lower panel shows combining anaphase A and B. Chromosomes are shown in blue, centrosomes in green, centrioles in black. Laterally associated microtubules are illustrated in orange. The end-on attached kinetochore microtubules (magenta in metaphase and green in anaphase) have the same length at both stages. The lower panel is an overlay of metaphase (magenta) with anaphase A (green, anaphase B movement was not considered) to show the relative movement of the autosomes with respect to the centrosomes. A rounding of the autosomes, a shrinking of the volume of the centrosomes and a change in the attachment angle of the microtubules is illustrated. (**B**) Tug-of-war model for the initiation of X chromosome segregation in anaphase I. The X chromosome (red), the holocentric kinetochore (gray), X chromosome-attached kinetochore microtubules (green). Left panel: At metaphase, pulling forces at the X chromosome are in balance. Right panel: segregation of the X chromosome is initiated by an imbalance of forces, obvious by an unequal number of kinetochore microtubules attached to the opposite sides of the X chromosome. X will move to the side with more attached kinetochore microtubules.

### Role of pulling forces during anaphase

The spermatocyte meiotic spindle shows two dramatic molecular distinctions during anaphase I that indicate pulling forces are critical to efficiently segregate chromosomes. First, outer kinetochore proteins are retained for the duration of both divisions. This is in contrast to mitosis and oocyte meiotic anaphase, where kinetochore levels diminish dramatically during anaphase progression (Asbury, 2017). Second, microtubules do not disassemble but instead remain attached to spermatocyte chromosomes, providing a stable connection to transmit the forces from poles needed to pull chromosomes apart. Indeed, we detected these end-on attached microtubules connections on the lagging X chromosome, which are subjected to pulling forces as poles move apart (**Fig. 5**).

Our results also suggest that *C. elegans* spermatocyte meiosis may not rely on a classical inter-chromosomal microtubule structure, or central spindle, for chromosome movement (Scholey et al., 2016). Acentrosomal oocyte meiosis (Dumont et al., 2010; Laband et al., 2017; Redemann et al., 2018; Yu et al., 2019) and centrosomal embryonic mitosis (Nahaboo et al., 2015; Yu et al., 2019), exhibit inter-chromosomal microtubules that push chromosomes apart. In contrast, in spermatocyte meiosis we see two distinct patterns. In XO males, the spindle midzone is dominated by a microtubule bridge connecting the unpaired X chromosome to the poles (**Fig. 1** and **6**). In XX hermaphrodites, *tra-2*(e1090) mutant males, or meiosis II, we observed few inter-chromosomal microtubules between the paired chromosomes (**Fig. S3B**). The absence of a classical inter-chromosomal microtubule structure may suggest a reliance on molecular components that generate pulling forces at the cellular cortex (Grill et al., 2003). We speculate this reliance may have evolved to help reliably resolve a lagging chromosome during anaphase I. Nonetheless, our results indicate that different cell types within a single organism distinctly regulate kinetochore and spindle structure to balance pulling and pushing of chromosomes in different contexts.

### Resolution of lagging chromosomes during sperm meiosis

Our studies reveal new features of the dynamics of chromosome lagging and resolution. We find continuous microtubules attach to each side of the lagging X chromosome throughout anaphase I. How do these microtubules lengthen? In one scenario, kinetochore microtubules could grow at their plus ends. As the spindle poles move apart, microtubule growth at a similar rate to spindle elongation could maintain association of kinetochore microtubules to the X chromosome. Alternatively, if the net growth rate of kinetochore microtubules exceeds the rate of spindle elongation, microtubules could attach laterally to the X, thus allowing minus-end directed interaction with motor proteins such as dynein (Reck-Peterson et al., 2018). Such a scenario could also explain the bending of microtubules we observed, possibly driven by dynein as it becomes processive towards the microtubule minus-end (Schmidt et al., 2005; Schmidt et al., 2017). As we find both end-on and laterally attached microtubules on the lagging X (**Fig. 7D**), their exact contributions to the segregation process remain to be determined.

A final question is how the lagging X chromosome resolves during anaphase I. We propose an imbalance of pulling forces that results stochastically via a continuous attachment and detachment of kinetochore microtubules. In such a ‘tug-of-war’ model, the side that maintains more connections to X wins, enabling resolution to one pole (**Fig. 11B**). Indeed, our EM and light microscopy data supports this model (**Fig. 8**). Such a tug-of-war mechanism has been suggested during chromosomal oscillations at mitotic prometaphase and metaphase (Ault et al., 1991; Skibbens et al., 1993; Soppina et al., 2009) with chromokinesins and dynein as possible candidates for switching the direction of the oscillations (Sutradhar and Paul, 2014).

*In toto*, our approach of combining cellular imaging within living males and quantification of 3D ultrastructure of staged spindles lays the groundwork for further studies on molecular mechanisms of chromosome segregation and provides analytical tools for biophysical analyses on spindles in a broad range of contexts (Fabig et al., 2016; Shakes et al., 2011; Winter et al., 2017). For example, many species have evolved distinct spindle and segregation strategies to resolve unequal numbers of sex chromosomes (Fabig et al., 2016). Recent work has also shown that segregation in cells with aneuploidy and chromosomal abnormalities are potential drivers of infertility (Barri et al., 2005; Garcia-Mengual et al., 2019; Hassold and Hunt, 2001; Ioannou and Tempest, 2015) and cancer progression (Bolhaqueiro et al., 2019; Chunduri and Storchova, 2019; Ly et al., 2019). Our studies can thus impact the understanding of partition mechanisms that can segregate both paired and lagging chromosomes to efficiently and reliably generate haploid sperm.

## Experimental procedures

### Strains and worm handling

#### Strains

The following strains were used in this study: N2 wild type (Brenner, 1974); MAS91 (unc-119(ed3) III; ItIs37[pAA64; pie-1::mCherry::HIS58]; ruIs57[pie-1::GFP::tubulin + unc-119(+)]) (Han et al., 2015); MAS96 (unc-119(ed3) III; ddIs6[tbg-1::GFP + unc-119(+)]; ltIs37[pAA64; pie-1::mCherry::HIS-58 + unc-119(+)] IV, qaIs3507[pie-1::GFP::LEM-2 + unc-119(+)]) (M. Srayko, Alberta);; TMR17 (unc-119(ed3) III; ddIs6[tbg-1::GFP + unc-119(+)]; ltIs37[pAA64; pie-1::mCherry::HIS-58 + unc-119(+)] IV) (this study); TMR18 (him-8(e1489) IV; unc-119(ed3) III; ddIs6[tbg-1::GFP + unc-119(+)]; ltIs37[pAA64; pie-1::mCherry::HIS-58 + unc-119(+)] IV) (this study); TMR26 (zim-2 (tm574) IV; unc-119(ed3) III; ddIs6[tbg-1::GFP + unc-119(+)]; ltIs37[pAA64; pie-1::mCherry::HIS-58 + unc-119(+)] IV) (this study); XC110 (tra-2(e1094)/dpy-10(e128) II; unc-119(ed3) III; ItIs37[pAA64; pie-1::mCherry::HIS58] (IV); ruIs57[pie-1::GFP::tubulin + unc-119(+)]) (this study); XC116 (tra-2(e1094)/dpy-10(e128) II; ddIs6[tbg-1::GFP + unc-119(+)]; ltIs37[pAA64; pie-1::mCherry::HIS-58 + unc-119(+)] IV) (this study); SP346 (tetraploid, 4n) (Madl and Herman, 1979).

#### Worm handling

Worms were grown on nematode growth medium (NGM) plates at 20°C with *E. coli* (OP50) as a food source (Brenner, 1974). Male worms were produced by exposing L4 hermaphrodites to 30°C for 4-6 h and checking the resulting progeny for male worms after three days (Sulston and Hodgkin, 1988). Males were maintained by mating 20-30 male worms with five L4 hermaphrodites. Triploid worms were obtained by mating tetraploid hermaphrodites with males of either MAS91 or TMR17. F1 male animals were selected and imaged as described below.

### Light microscopy and analysis of spindle dynamics

#### Light microscopy

Age-synchronized males (3 d after bleaching adult hermaphrodites fertilized by males) were placed in droplets of 1 µl polystyrene microbeads solution (bead diameter of 0.1 µm; Polysciences, USA) on 10% agarose pads. Samples were then covered with a coverslip and sealed with wax (Kim et al., 2013). For a comparative analysis of mitotic and female meiotic embryos, 3 d old hermaphrodites were dissected in M9 buffer and transferred to 4% agarose pads. We used a confocal spinning disk microscope (IX 83, Olympus, Japan) equipped with a 60x 1.2 NA water immersion objective and an EMCCD camera (iXon Ultra 897, Andor, UK) for live-cell imaging. The meiotic region within single males was imaged for about one hour and a z-stack was recorded either every 20 s or 30 s. Z-stacks for embryos were recorded every 20 s for about 1 h. Images were then corrected for photobleaching using the Fiji software package (Schindelin et al., 2012).

#### Analysis of spindle dynamics

Image stacks were analyzed with the Arivis Vision4D software package (arivis AG, Germany). Individual spindles were cropped and spindle poles in each frame were segmented by thresholding. The Euclidean distance of the center of mass of both spindle poles was then calculated for each time point. For the production of kymographs, the original image data was resampled with a custom-made python script in arivis Vision4D. The spindle axes were rotated in all three dimensions to align the axis along the z-direction. As a consequence each spindle had a comparable orientation with an isotropic voxel size of 0.1 µm and a radius of 0.9 µm around the spindle axis. All voxels were then recalculated based on the initial transformation of the axis with an extrapolation of 1 µm at each pole in the direction of the axis. As the axes of the spindles were chosen to lay in the z-dimension all images in the resampled datasets were laying orthogonally to it (x, y-plane). For the calculation of kymographs, the Gaussian weighted sum of fluorescence in each plane was calculated along the spindle axis and repeated for all time points. For the analysis of chromosome movements, the peak maxima of the chromosome fluorescence signals were then used to calculate the distances for each time point. Individual measurements were aligned according to the onset of anaphase and the mean distance was then calculated and plotted against time relative to anaphase onset. For characterizing the dynamic properties of spindles these mean values were then used to determine spindle length at metaphase and after anaphase. The initial speed of spindle elongation and chromosome movement was calculated by fitting a linear function to the measurements during the first minute after anaphase onset as the segregation speed slowed down continuously.

To illustrate the process of division, the spindles were resampled and rotated as described above but with a radius of 3 µm around the spindle axis and an extrapolation of 2 µm after the spindle poles. Then an y,z-projection over x (maximum intensity) was calculated for each time point to display the resampled volume as a plane image (**Fig. 1-3**). For a comparison of microtubule density on both sides of the X chromosome facing the spindle poles, the sum of fluorescence was calculated within two cubic boxes (with a similar volume of 1 µm³) adjacent to the X chromosome in the resampled light microscopic image data. The box on the side, where the chromosome moved to at the time of segregation, was termed “volume 1”, the other “volume 2”. The ratio between both values at each time point indirectly describes the difference in the number of microtubules (**Fig. 8E-F**).

For each data set, the visco-elastic property of the X chromosome was probed by segmenting it in a resampled 3D dataset and measuring its dimensions. Along the spindle axis, the length of the X chromosome was measured (z-dimension). Orthogonal to the x-axis, the mean values for the x- and y-dimension were calculated. A shape coefficient was then calculated (z/[(x+y)/2]) to illustrate the change of the shape of the X chromosome over time (**Fig. 5A-B**).

The centrosomes were segmented in 3D image data from worms expressing γ-tubulin::GFP and histone H2B::mCherry with the Arivis Vision4D software package by applying a cut-off threshold to the 3D image data. All fluorescence signals above the threshold were included in the segment of the centrosomes. The volume of the segments was then calculated for each frame and each centrosome individually for spindles in meiosis I and II. When centrosomes split in meiosis I and could be segmented individually both volumes were summed together for the respective frame (**Fig. S1**).

### Immunostaining for light microscopy

For antibody staining of *C. elegans* gonads, synchronized males were dissected and fixed in 1% paraformaldehyde using established protocols (Howe et al., 2001). Methanol/acetone fixation was used for immunolabeling of mitotic and meiotic embryos (Shakes et al., 2009). Primary and secondary antibodies were diluted in blocking buffer (PBS + 0.1% Tween 20 and 10 mg/ml BSA) and staining was conducted at room temperature in a humid chamber. Primary antibodies were used in overnight incubations (unless otherwise noted). Commercial sources or labs kindly providing antibodies were as listed: 1:200 mouse anti-NDC-80 (Novus Biologicals, catalog #42000002); 1:200 mouse anti-α-tubulin (DM1A Sigma-Aldrich, catalog #T6199); 1:500 rabbit anti-KNL-1 (Desai et al., 2003); 1:500 rabbit anti-KNL-3 (Cheeseman et al., 2005); 1:400 rabbit anti-CENP-C^HCP-4^ (Moore et al., 2005); 1:400 rabbit anti-HIM-10 (Skop et al., 2001); and 1:50 FITC-conjugated anti-α-tubulin (Sigma-Aldrich, #F2168). Secondary antibodies included: goat anti-rabbit AlexaFluor 488-labeled IgG (used at 1:200); goat anti-mouse AlexaFluor 488-labeled IgG (used at 1:200); goat anti-mouse AlexaFluor 564-labeled IgG (used at 1:200); and donkey anti-rabbit Cy3 (used at 1:500). DNA was visualized using DAPI at 0.1 µg/ml. Slides were prepared by using VectaShield (Vector Labs, USA) as a combined mounting and anti-fade medium. Confocal images were acquired using a Zeiss LSM710 microscope with Zen software (**Fig. 4**). Super-resolution images were collected using an OMX 3D-SIM microscope (GE Healthcare, USA) with an Olympus (Shinjuku, Japan) 100x UPlanSApo 1.4 NA objective (Olympus, Japan). Images were captured in z-steps of 0.125 μm and processed using SoftWoRx (GE Healthcare, USA) and IMARIS (Bitplane, Switzerland) 3D imaging software (**Fig. S4**).

### Laser microsurgery

Age-synchronized males (3d old) were placed within a of droplet of 1 µl M9 buffer containing 1mM levamisole and 0.1 µm polystyrene microbeads (Polysciences, USA) on a 10 % agarose pad. Samples were then covered with a coverslip and sealed with wax. For imaging during laser microsurgery, we used a confocal spinning disk microscope (Ti Eclipse, Nikon, Japan) equipped with a 60x 1.2 NA water immersion objective, a 1.5x optovar, an EMCCD camera (iXon Ultra 897, Andor, UK) and a mode-locked femtosecond Ti:sapphire laser (Chameleon Vision II, Coherent, USA) operated at a wavelength of 800 nm. After locating spindles in anaphase I within males, a single image was recorded in intervals of 1 s. Subsequently, a position for the laser cut was chosen and a single spot with a diameter of about 1.3 µm was ablated with a laser power of 150 mW and an exposure time of 30 ms. Image acquisition was continued until the X chromosome had been fully segregated (**Fig. 5C-D**). For further analysis the images were corrected for photobleaching within the Fiji software package and corrected for movement using the plugin “image stabilizer” (http://www.cs.cmu.edu/~kangli/code/Image_Stabilizer.html; February 2008).

### Specimen preparation for electron microscopy

Males were ultra-rapidly frozen using a HPF COMPACT 01 high-pressure freezer (Engineering Office M. Wohlwend, Sennwald, Switzerland). For each freezing run, five individuals were placed in a type-A aluminum planchette (100 µm deep; Wohlwend, article #241) pre-wetted with hexadecene (Merck) and then filled with M9 buffer containing 20% (w/v) BSA (Roth, Germany). The specimen holders were closed by gently placing a type-B aluminum planchette (Wohlwend, article #242) with the flat side facing the sample on top of a type-A specimen holder. The sandwiches were frozen under high pressure (∼2000 bar) with a cooling rate of ∼20000°C/s (Fabig et al., 2019). Specimen holders were opened under liquid nitrogen and transferred to cryo-vials filled with anhydrous acetone containing 1% (w/v) osmium tetroxide (EMS) and 0.1% (w/v) uranyl acetate (Polysciences, USA). Freeze substitution was performed in a Leica AFS (Leica Microsystems, Austria). Samples were kept at −90°C, then warmed up to −30°C with steps of 5°C/h, kept for 5 h at −30°C and warmed up again (steps of 5°C/h) to 0°C. Subsequently, samples were washed three times with pure anhydrous acetone and infiltrated with Epon/Araldite (EMS, USA) epoxy resin at increasing concentrations of resin (resin:acetone: 1:3, 1:1, 3:1, then pure resin) for 2h each step at room temperature (Muller-Reichert et al., 2003). Samples were incubated with pure resin over night and then for 4 h. For thin-layer embedding samples were placed between two Teflon-coated glass slides and allowed to polymerize at 60°C for 48 h (Muller-Reichert et al., 2008). Polymerized samples were remounted on dummy blocks and semi-thin serial sections (300 nm) were cut using an EM UC6 (Leica Microsystems, Austria) ultramicrotome. Ribbons of sections were collected on Formvar-coated copper slot grids, post-stained with 2% (w/v) uranyl acetate in 70% (v/v) methanol and 0.4% (w/v) lead citrate and allowed to dry prior to inspection.

### Electron tomography, microtubule segmentation and stitching of data sets

In preparation for electron tomography, both sides of the samples were coated with 15 nm-colloidal gold (BBI, UK). To select cells in meiosis, serial sections were pre-inspected at low magnification (∼2900x) using a Zeiss EM906 transmission electron microscope (Zeiss, Germany) operated at 80 kV. Serial sections containing cells/regions of interest were then transferred to a Tecnai F30 transmission electron microscope (ThermoFischer Scientific, USA) operated at 300 kV and equipped with a US1000 CCD camera (Gatan, USA). Tilt series were acquired from −65° to +65° with 1° increments at a magnification of 4700x. Specimens were then rotated 90° to acquire a second tilt series for double-tilt electron tomography (Mastronarde, 1997). Electron tomograms were calculated using the IMOD software package (Kremer et al., 1996). As previously described (Redemann et al., 2014; Weber et al., 2012), microtubules were automatically segmented using the ZIBAmira (Zuse Institute Berlin, Germany) software package (Stalling et al., 2005).

Individual tomograms were then stitched and combined (Weber et al., 2014) to represent whole microtubule networks in 3D models (Redemann et al., 2017). Chromosomes, kinetochores and centrioles were manually segmented. Kinetochores were modeled around each chromosome by gradually increasing the chromosome volume until the area of the ribosome-free zone around each chromosome (Howe et al., 2001; O’Toole et al., 2003) was covered, giving a thickness of the male meiotic holocentric kinetochore of about 150 nm.

### Analysis of tomographic data

#### Classification of microtubules

First, the distance between each point of a microtubule segment and the closest point of the surface of individual chromosomes was calculated. Only microtubules within a distance of 150 nm or less were considered kinetochore microtubules as this distance was measured to be the approximate extent of the kinetochore in the electron tomograms. The kinetochore is visible in the electron tomograms as a less stained region around the chromosomes (Howe et al., 2001). Additionally, each kinetochore microtubule was assigned to the X chromosome or to one of the autosomal chromosomes according to its closest distance to the chromosome surface. As microtubules in anaphase pass between the autosomes and attach to the X chromosome after that, they were first checked for an interaction with the X chromosome and if there was none, further analysis was performed to check for a potential autosomal interaction. For each chromosome the microtubule interactions were subdivided between end-on and lateral. We defined an end-on association by extrapolating the microtubule after its end for 150 nm and checking if this extrapolated line was cutting the surface of the chromosome. If that criterion was not met, we considered the association of the microtubule with the given chromosome as lateral.

#### Length distribution

Furthermore, we analyzed the length distribution of microtubules. The length distribution for each microtubule class in each meiotic spindle is presented as a violin plots in order to assess the mean, standard deviation as well as the distribution of the data. For that, individual data points were binned into 25 intervals and the width of the plot was normalized to equalize relative densities within the individual datasets. Further, the mean and the standard deviation were shown and the variance among the datasets was compared using a one-way analysis of variance (ANOVA; **Fig. 10, Fig. S6, Fig. S8**).

We also analyzed the ratio of the sum of microtubule length between two defined volumes analogous to the analysis of the light microscopic data. For that a box of 1 µm³ was placed on either side of the X chromosome facing the spindle poles. The microtubules within this box were extracted and their length was measured and summed up. The ratio of the box closer to the respective pole against the second box was calculated (**Fig. 8A-B**). The microtubule tortuosity (microtubule spline length divided by end-end length; **Fig. 7C-D**) was measure for end-on and lateral microtubules in contact with the X chromosome.

#### Chromosome shape

Further, we analyzed the shape of the chromosomes in the EM data as previously described (Lindow et al., 2018). In brief, chromosomes were manually segmented and along the pole-to-pole axis of the spindle orthogonal planes were placed with 10 nm spacing. For every plane the area was calculated that intersects the individual chromosome surface. After plotting the area against the pole-pole distance a Gaussian function containing five terms was fit with MATLAB (MATLAB 2017b, The MathWorks, USA) and the full width at half maximum (FWHM) for each chromosome was determined and compared (**Fig. 10A-B**).

For measuring the distance between centrioles and the end-on microtubule end at the autosomes, we first selected the closest centriole at the putative microtubule minus-end. Then we extracted the position of the respective putative plus-end and calculated the Euclidean distance between the centriole and the putative plus-end (**Fig. 10C-D**).

The angle between the microtubule plus-end and the chromosome-centrosome axis was determined by calculating the vector between the respective chromosome and the centrosome and the vector between chromosome and the respective end. Then the angle between both vectors was calculated (**Fig. 10E-F**).

## Acknowledgements

The authors would like to thank Dr. Michael Laue (RKI, Berlin, Germany) for using the COMPACT 01 (Wohlwend) high-pressure freezer, the Core Facility Cellular Imaging of the Faculty of Medicine Carl Gustav Carus (TU Dresden, Germany) and the light- and electron microscopy facilities at the MPI-CBG (Dresden, Germany) for technical assistance. We are also grateful to Drs. Diane Shakes (Williamsburg VA, USA), Stefanie Redemann (Charlottesville VA, USA) and Kevin O’Connell (Bethesda MD, USA) for a critical reading of the manuscript. We would like to thank Martin Merkel, Ewa Kania and Sophia May for help in tomographic reconstruction and microtubule segmentation. The authors are grateful to Falko Löffler, Carola Bender and Christian Götze (Arivis AG) for help with image processing in arivis Vision4D. Some strains were provided by the CGC, which is funded by NIH Office of Research Infrastructure Programs (P40 OD010440). We acknowledge NIH grant NIH1S10OD024988-01 for the purchase on the OMX microscope. Work in the Müller-Reichert laboratory is supported by funds from the Deutsche Forschungsgemeinschaft (MU 1423/10-1). R.K. received funding from the European Union’s Horizon 2020 research and innovation program under the Marie Skłodowska-Curie grant agreement No. 675737 (grant to T.M.R.). Work in the Chu lab is supported by the NIH grant R03 HD093990-01A1 and the NSF Awards RUI-1817611 and DBI-1548297.

## Supplementary Figures

**Figure S1.**
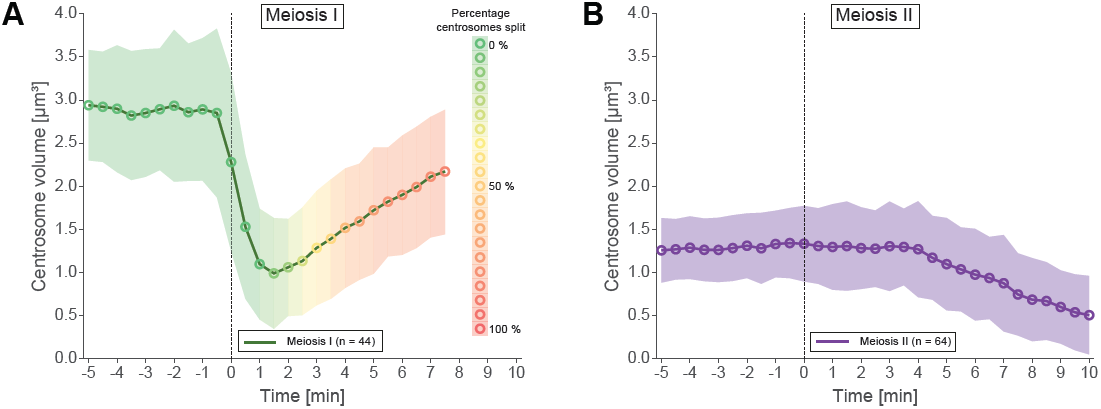
Analysis of centrosomal volumes in meiosis I and II. (**A**) Plot showing centrosome volume in meiosis I over time. The 3D volume was measured in worms expressing γ-tubulin::GFP and histone H2B::mCherry. For each dataset a fixed threshold was defined to segment the outer border of the centrosome. The mean volume is plotted as a green line for unsplitted centrosomes and shown as an orange to red line for splitted centrosomes. The percentage of splitted centrosomes is indicated by this color change. For splitted centrosomes, the sum of both separated centrosomes was determined (n=44). The standard deviation is depicted as a shaded area. (**B**) Centrosome volume over time (purple line) in meiosis II (n=64).

**Figure S2.**
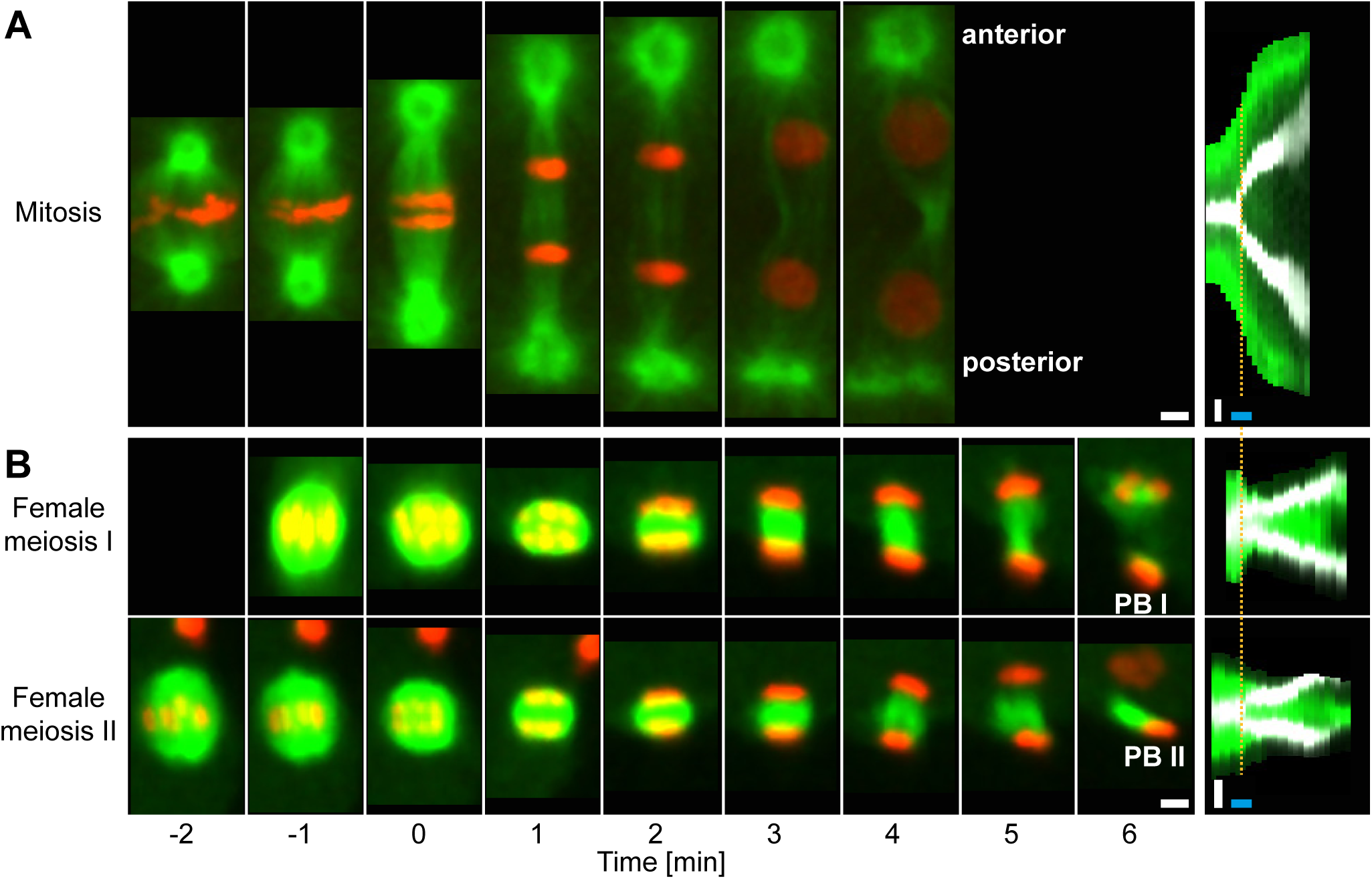
Spindle organization in early embryos and oocytes of *C. elegans*. (**A**) First mitosis in the early embryo. Microtubules (β-tubulin::GFP, green) and chromosomes (histone H2B::mCherry, red) are visualized in time series of confocal image projections (left panel). Anaphase onset is time point zero (t=0). The anterior and the posterior centrosome is indicated). The progression of chromosome segregation is shown in a kymograph (right panel). Anaphase onset is indicated by a dashed line (orange). The histone H2B::mCherry signal in the kymograph is shown in gray. Scale bar (white), 2 µm; time bar (blue), 2 min. (**B**) Female meiosis in hermaphrodites. The two consecutive meiotic divisions show acentrosomal spindles (left panels; upper row, meiosis I; lower row, meiosis II). Chromosomes are extruded by two polar bodies (PB I and II). Kymographs (right panels) illustrate the reorganization of the microtubules (left panels). Scale bar (white), 2 µm; time bar (blue), 2 min.

**Figure S3.**
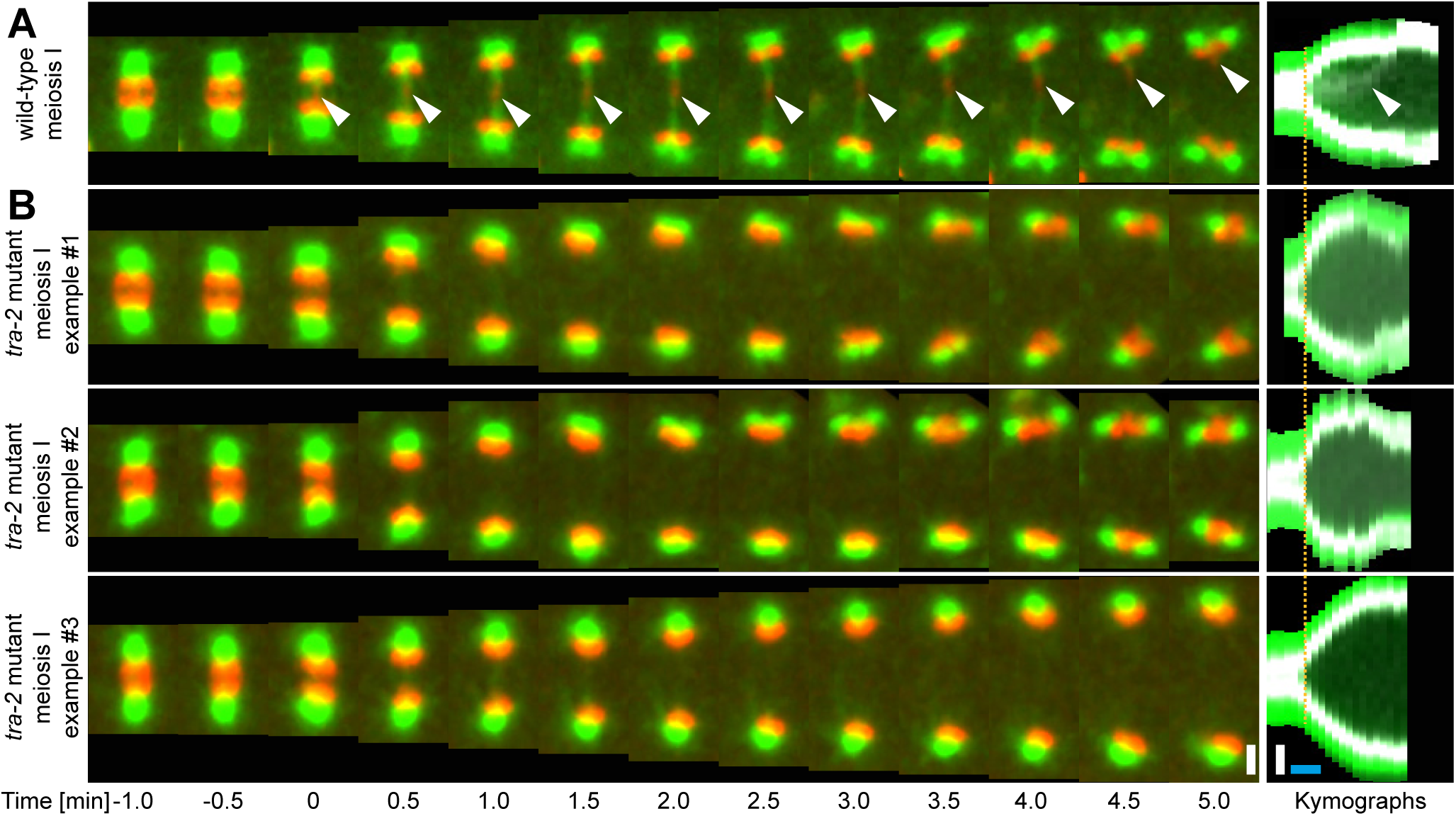
Spindle organization in meiosis I in wild-type and *tra-2* mutants. Time series of confocal image projections of meiosis I spindles in **(A)** wild-type and in **(B)** three examples of *tra-2*(e1094) mutant males. Microtubules (β-tubulin::GFP) and chromosomes (histone H2B::mCherry) are visualized in green and red, respectively. Anaphase onset is time point zero (t=0). Chromosome segregation is also visualized in kymographs (right panel; the start of the separation of the autosomes is indicated by an orange line). Scale bars (white), 2 µm; time bar (blue), 2 min.

**Figure S4.**
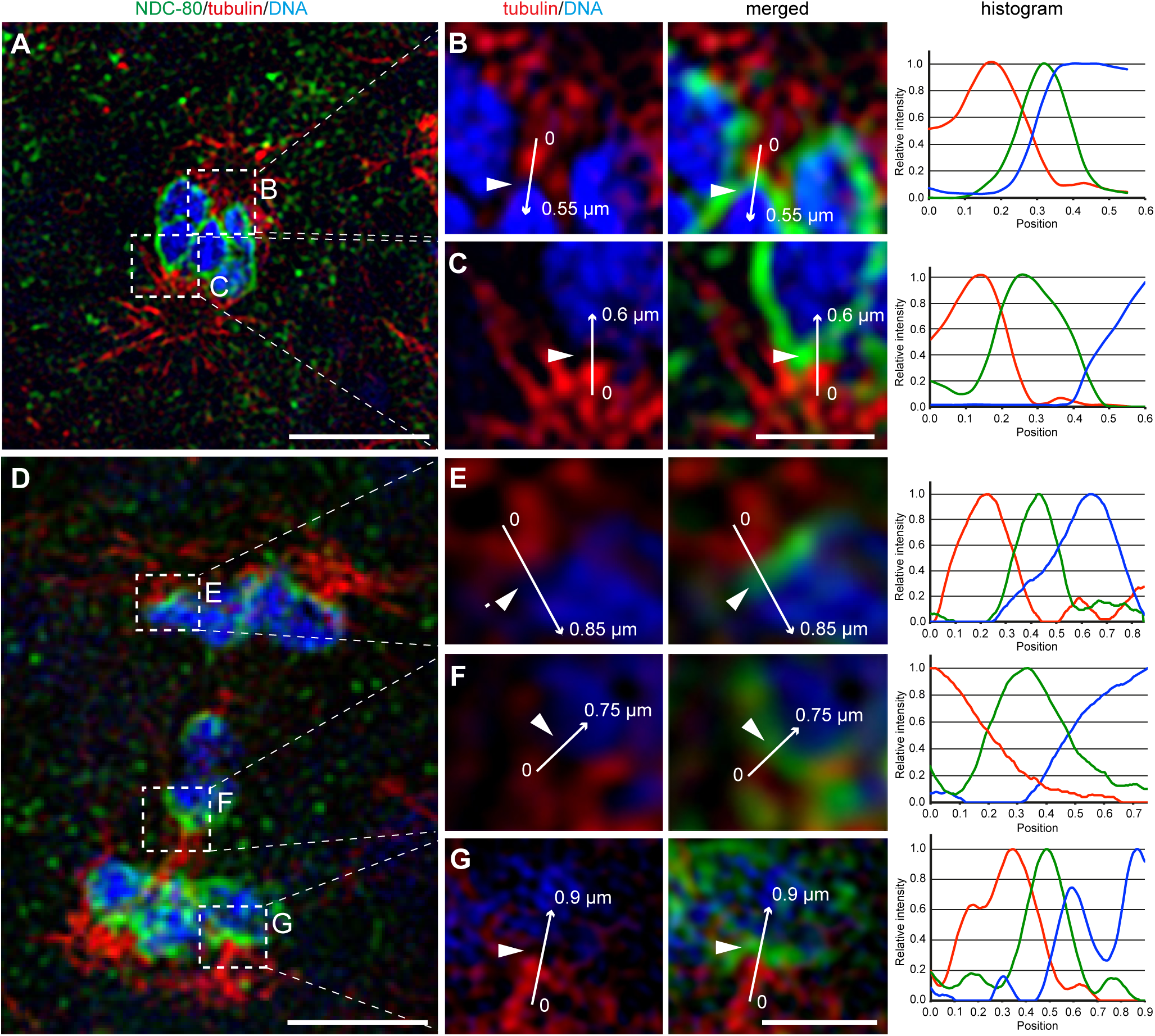
The outer kinetochore protein NDC80 localizes between chromosomes and microtubules at spermatocyte metaphase and anaphase I. (**A**) Super-resolution fluorescence microscopy of metaphase I in fixed *him-8* X0 males stained with antibodies against α-tubulin (red) and NCD-80 (green). DAPI stained DNA is in blue. Scale bar, 2 µm. (**B-C**) Enlargement of boxed regions as shown in (A) highlighting microtubule and NDC-80 localization relative to metaphase chromosomes. Normalized intensity values along the arrows for each staining pattern are plotted in the histograms (right panels). Scale bars, 0.5 µm. (**D**) Super-resolution fluorescence microscopy of anaphase I in *him-8* X0 males. Imaging conditions were as given in (A). Scale bar, 2 µm. (**E-G**) Enlargement of boxed regions as shown in (A) highlighting microtubule and NDC-80 localization relative to separating chromosomes. Scale bar, 0.5 µm.

**Figure S5.**
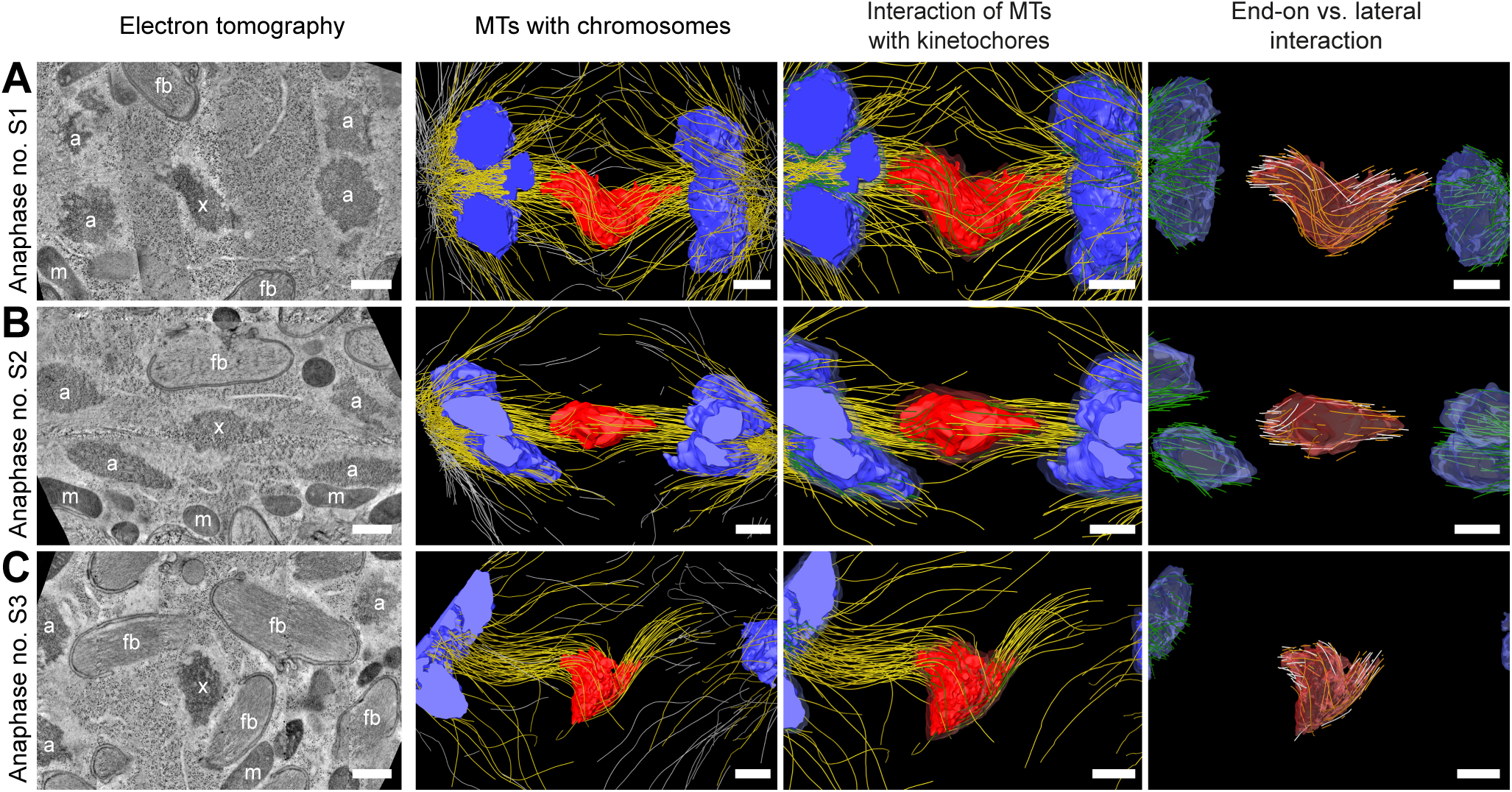
Visualization of partially reconstructed spindles in mid/late anaphase I. (**A**) Tomographic reconstruction of an anaphase I spindle. Left panel: tomographic slice showing the autosomes (a), and the univalent X chromosome (x) aligned along the spindle axis. Mitochondria (m) and fibrous body-membranous organelles (fb) are also indicated. Mid left panel: corresponding three-dimensional model illustrating the organization of the full spindle. Autosomes are in blue, the X chromosome in red, microtubules within a distance of 150 nm or closer to the chromosome surfaces in yellow and all other microtubules in gray. Mid right panel: association of microtubules with the kinetochores. Kinetochores are shown as semi-transparent regions around each chromosome. The part of each microtubule entering the kinetochore region around the holocentric chromosomes is in green. Right panel: visualization of end-on (white) *versus* lateral (orange) association of microtubules with chromosomes. Only the part inside of the kinetochore is shown. (**B**) Second example of spindle organization at mid anaphase I. (**C**) Example of spindle organization at late anaphase I. Scale bars, 500 nm.

**Figure S6.**
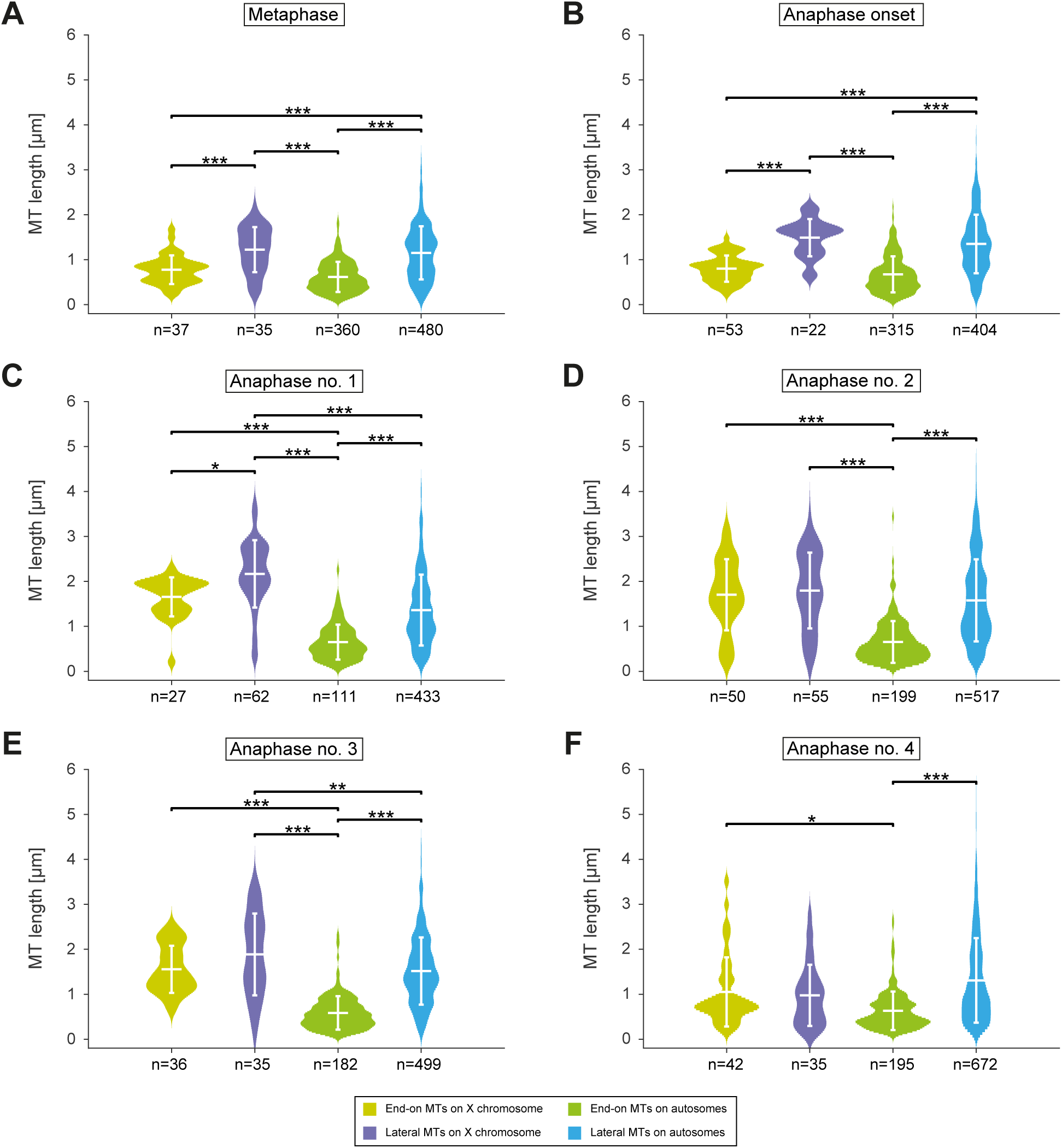
Length distribution of microtubules at different stages of meiosis I. (**A**) Analysis of microtubule length in metaphase (corresponding to data sets as shown in Fig. 6). Measurements for: end-on X chromosome-attached microtubules (yellow), lateral X chromosome-attached microtubules (violet), end-on autosome-attached microtubules (green), and lateral autosome-attached microtubules (light blue). The mean and standard deviation are shown as white lines in the violin plots. Distributions were compared using a one-way ANOVA with three levels of significance: * is p <= 0.05; ** is p <= 0.01; and *** is p <= 0.001. (**B**) Microtubule length at anaphase onset. (**C**) Microtubule length at mid anaphase (dataset anaphase 1 of Fig. 6). (**D**) Microtubule length at mid anaphase (dataset anaphase 2 of Fig. 6). (**E**) Microtubule length at mid anaphase (dataset anaphase 3 of Fig. 6). (**F**) Microtubule length at late anaphase (dataset anaphase 4 of Fig. 6).

**Figure S7.**
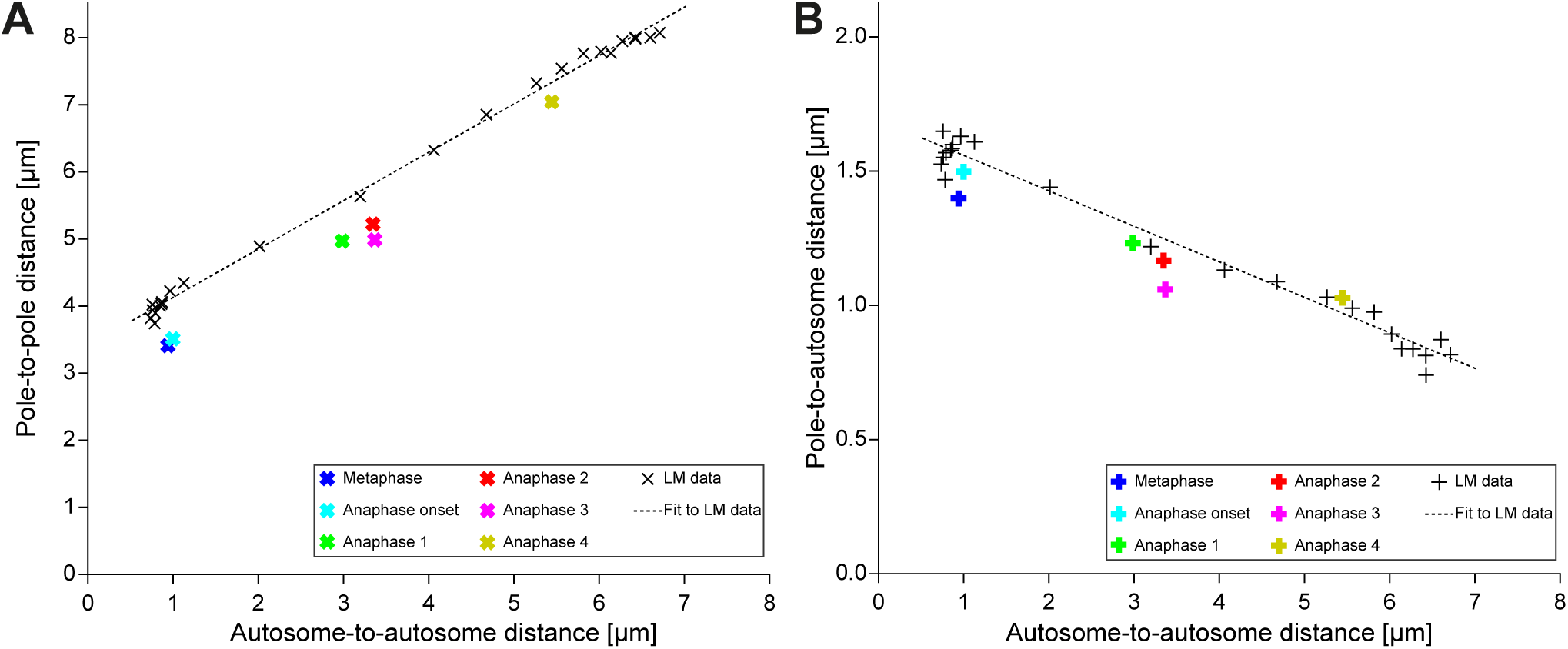
Comparison of the EM data sets with the dynamic light microscopic data. (**A**) The pole-to-pole distance plotted against the autosome-to-autosome distance for each data set (stages and color coding as presented in Fig. 8B). The colored bold “x” symbols illustrate where the EM-data sets are positioned with respect to the averaged light microscopy data as shown in Fig. 3E (black “x” symbols with fitted black dashed line). (**B**) The pole-to-autosome distance plotted against the autosome-to-autosome distance in meiosis I (bold-colored “+” symbols show the EM data, black symbols and dashed line illustrate the light microscopic data with linear fit).

**Figure S8.**
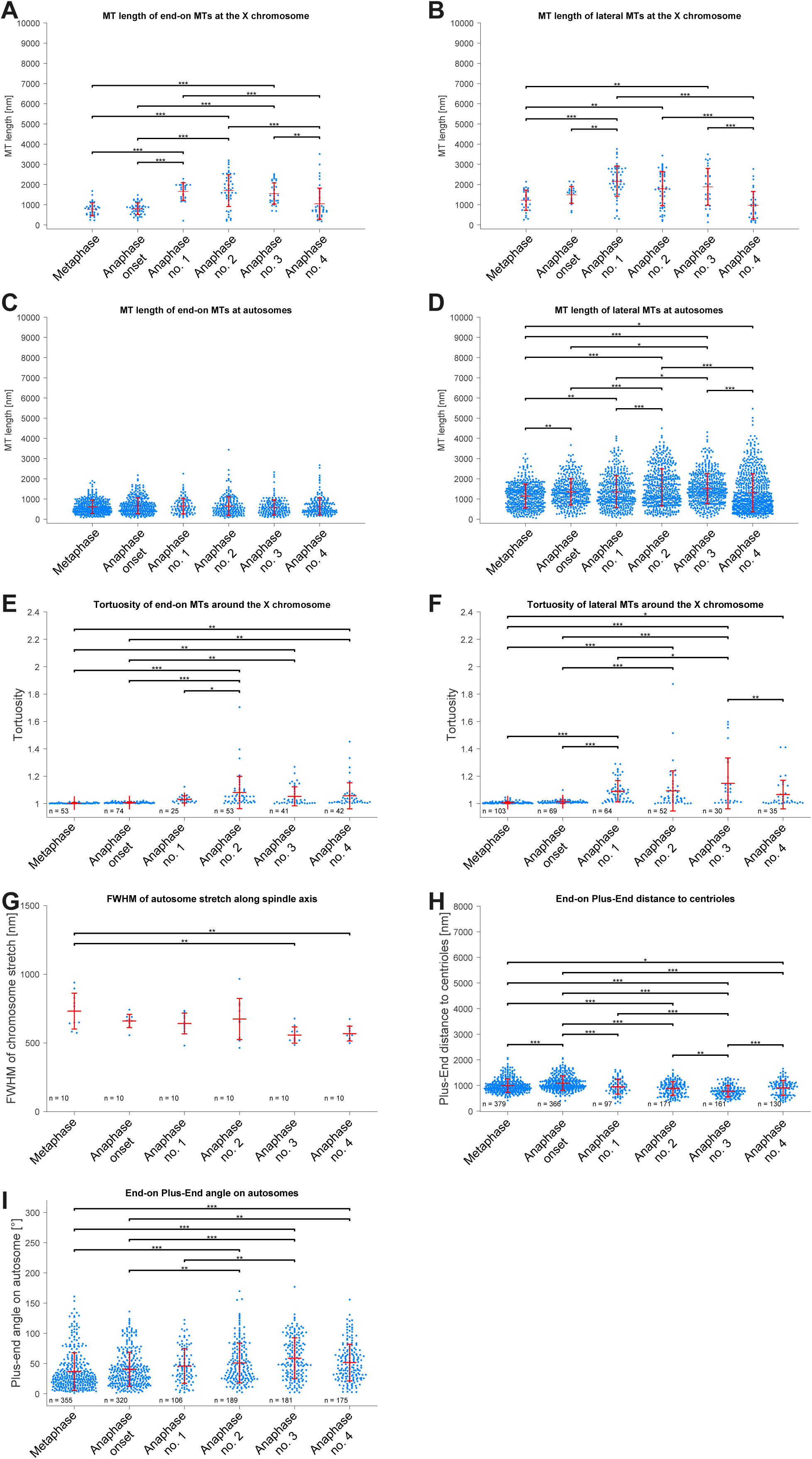
Comparison of distributions within EM data measurements. (**A**) The length of end-on kinetochore microtubules associated with the X chromosome for each meiotic stage corresponding to Fig. 7A. The mean, standard deviation and single measurements for each meiotic stage (according to the ones as shown in Figure 6) are given. Results of a one-way analysis of variance (ANOVA) of all data sets against each other are shown. Level of significance: * is p <= 0.05; ** is p <= 0.01; and *** is p <= 0.001. (**B**) Plot showing the length of lateral kinetochore microtubules associated with the X chromosome for each meiotic stage corresponding to Fig. 7B. (**C**) The length of end-on kinetochore microtubules associated with the autosomes for each meiotic stage corresponding to Fig. 9A. (**D**) The length of lateral s associated with the autosomes for each meiotic stage corresponding to Fig. 9B. (**E**) The tortuosity of end-on kinetochore microtubules associated with the X chromosome for each meiotic stage corresponding to Fig. 7D. (**F**) The tortuosity of lateral kinetochore microtubules associated with the X chromosome for each meiotic stage corresponding to Fig. 7D. (**G**) The FWHM of the chromosome stretch for each meiotic stage corresponding to Fig. 10B. (**H**) The distance of kinetochore microtubule plus-ends at the autosomes to centrioles for each meiotic stage corresponding to Fig. 10D. (**I**) The angle measurements as given in Fig. 10E for each meiotic stage corresponding to Fig. 10F.

## Supplementary Movies

Movie S1. **Live-cell imaging of the first and second meiotic division in wild-type males.**

The strain was labeled with β-tubulin::GFP (green) and histone H2B::mCherry (red) to visualize microtubules and chromosomes, respectively. Time is given relative to anaphase onset of meiosis I. Scale bar, 2 µm. This supplementary movie corresponds to Fig. 1A and B.

Movie S2. **Live-cell imaging of the first meiotic division in wild-type males.**

The strain was labeled with γ-tubulin::GFP (green) and histone H2B::mCherry (red) to visualize centrosomes and chromosomes, respectively. The image data was resampled to correct for the movements of the male worm. Time is given relative to anaphase onset. Scale bar, 2 µm. This supplementary movie corresponds to Fig. 3B and Fig. S1A.

Movie S3. **Live-cell imaging of the second meiotic division in wild-type males.**

The strain was labeled with γ-tubulin::GFP (green) and histone H2B::mCherry (red) to visualize centrosomes and chromosomes, respectively. The image data was resampled to correct for the movements of the male worm. Time is given relative to anaphase onset. Scale bar, 2 µm. This supplementary movie corresponds to Fig. 3D and Fig. S1B.

Movie S4. **Laser ablation of the microtubule bridge in a meiotic spermatocyte spindle undergoing the first division.**

Spindle within an immobilized wild-type male worm labeled with β-tubulin::GFP (green) and histone H2B::mCherry (red) to visualize microtubules and chromosomes, respectively. The applied single laser cut is indicated by a circle. Time is given relative to the laser cut. Scale bar, 2 µm. This supplementary movie corresponds to Fig. 5C (experiment no. 1).

Movie S5. **Double-cut laser ablation of the microtubule bridge in a meiotic spermatocyte spindle undergoing first meiotic division.**

Spindle within an immobilized wild-type male worm labeled with β-tubulin::GFP (green) and histone H2B::mCherry (red) to visualize microtubules and chromosomes, respectively. The laser cuts are indicated by circles. Time is given relative to the first laser cut. Scale bar, 2 µm. This supplementary movie corresponds to Fig. 5D.

Movie S6. **Full tomographic reconstruction of the metaphase I spindle in a wild-type male spermatocyte.**

Tomographic slices through the entire 3D volume and corresponding three-dimensional model illustrating the organization of the full spindle. Autosomes are shown in different grades of either blue or cyan, the X chromosome in red, centriolar microtubules in purple, microtubules within a distance of 150 nm or closer to the chromosome surfaces in yellow and all other microtubules in gray. This supplementary movie corresponds to Fig. 6A.

Movie S7. **Full tomographic reconstruction of the spindle at the onset of anaphase I in a wild-type male spermatocyte.**

Three-dimensional model illustrating the organization of the full spindle. Autosomes are shown in different grades of either blue or cyan, the X chromosome in red, centriolar microtubules in purple, microtubules within a distance of 150 nm or closer to the chromosome surfaces in yellow and all other microtubules in gray. This supplementary movie corresponds to Fig. 6B.

Movie S8. **Full tomographic reconstruction of the anaphase I spindle (anaphase no. 1) in a wild-type male spermatocyte.**

Three-dimensional model illustrating the organization of the full spindle. Autosomes are shown in different grades of either blue or cyan, the X chromosome in red, centriolar microtubules in purple, microtubules within a distance of 150 nm or closer to the chromosome surfaces in yellow and all other microtubules in gray. This supplementary movie corresponds to Fig. 6C.

Movie S9. **Full tomographic reconstruction of the anaphase I spindle (anaphase no. 2) in a wild-type male spermatocyte.**

Three-dimensional model illustrating the organization of the full spindle. Autosomes are shown in different grades of either blue or cyan, the X chromosome in red, centriolar microtubules in purple, microtubules within a distance of 150 nm or closer to the chromosome surfaces in yellow and all other microtubules in gray. This supplementary movie corresponds to Fig. 6D.

Movie S10. **Full tomographic reconstruction of the anaphase I spindle (anaphase no. 3) in a wild-type male spermatocyte.**

Tomographic slices through the entire 3D volume and corresponding three-dimensional model illustrating the organization of the full spindle. Autosomes are shown in different grades of either blue or cyan, the X chromosome in red, centriolar microtubules in purple, microtubules within a distance of 150 nm or closer to the chromosome surfaces in yellow and all other microtubules in gray. This supplementary movie corresponds to Fig. 6E.

Movie S11. **Full tomographic reconstruction of the anaphase I spindle (anaphase no. 4) in a wild-type male spermatocyte.**

Three-dimensional model illustrating the organization of the full spindle. Autosomes are shown in different grades of either blue or cyan, the X chromosome in red, centriolar microtubules in purple, microtubules within a distance of 150 nm or closer to the chromosome surfaces in yellow and all other microtubules in gray. This supplementary movie corresponds to Fig. 6F.

## References

Albertson, D.G. 1984. Formation of the first cleavage spindle in nematode embryos. Dev Biol. 101:61–72.

Albertson, D.G., and J.N. Thomson. 1993. Segregation of holocentric chromosomes at meiosis in the nematode, Caenorhabditis elegans. Chromosome Res. 1:15–26.

Asbury, C.L. 2017. Anaphase A: Disassembling Microtubules Move Chromosomes toward Spindle Poles. Biology (Basel). 6.

Ault, J.G., A.J. DeMarco, E.D. Salmon, and C.L. Rieder. 1991. Studies on the ejection properties of asters: astral microtubule turnover influences the oscillatory behavior and positioning of mono-oriented chromosomes. J Cell Sci. 99 (Pt 4):701–710.

Barri, P.N., J.M. Vendrell, F. Martinez, B. Coroleu, B. Aran, and A. Veiga. 2005. Influence of spermatogenic profile and meiotic abnormalities on reproductive outcome of infertile patients. Reprod Biomed Online. 10:735–739.

Bennabi, I., M.E. Terret, and M.H. Verlhac. 2016. Meiotic spindle assembly and chromosome segregation in oocytes. J Cell Biol. 215:611–619.

Bolhaqueiro, A.C.F., B. Ponsioen, B. Bakker, S.J. Klaasen, E. Kucukkose, R.H. van Jaarsveld, J. Vivie, I. Verlaan-Klink, N. Hami, D.C.J. Spierings, N. Sasaki, D. Dutta, S.F. Boj, R.G.J. Vries, P.M. Lansdorp, M. van de Wetering, A. van Oudenaarden, H. Clevers, O. Kranenburg, F. Foijer, H.J.G. Snippert, and G. Kops. 2019. Ongoing chromosomal instability and karyotype evolution in human colorectal cancer organoids. Nat Genet. 51:824–834.

Brenner, S. 1974. The genetics of Caenorhabditis elegans. Genetics. 77:71–94.

Cheerambathur, D.K., B. Prevo, N. Hattersley, L. Lewellyn, K.D. Corbett, K. Oegema, and A. Desai. 2017. Dephosphorylation of the Ndc80 Tail Stabilizes Kinetochore-Microtubule Attachments via the Ska Complex. Dev Cell. 41:424–437 e424.

Cheeseman, I.M., J.S. Chappie, E.M. Wilson-Kubalek, and A. Desai. 2006. The conserved KMN network constitutes the core microtubule-binding site of the kinetochore. Cell. 127:983–997.

Cheeseman, I.M., I. MacLeod, J.R. Yates, 3rd, K. Oegema, and A. Desai. 2005. The CENP-F-like proteins HCP-1 and HCP-2 target CLASP to kinetochores to mediate chromosome segregation. Current biology : CB. 15:771–777.

Chunduri, N.K., and Z. Storchova. 2019. The diverse consequences of aneuploidy. Nat Cell Biol. 21:54–62.

Ciferri, C., S. Pasqualato, E. Screpanti, G. Varetti, S. Santaguida, G. Dos Reis, A. Maiolica, J. Polka, J.G. De Luca, P. De Wulf, M. Salek, J. Rappsilber, C.A. Moores, E.D. Salmon, and A. Musacchio. 2008. Implications for kinetochore-microtubule attachment from the structure of an engineered Ndc80 complex. Cell. 133:427–439.

Crowder, M.E., M. Strzelecka, J.D. Wilbur, M.C. Good, G. von Dassow, and R. Heald. 2015. A comparative analysis of spindle morphometrics across metazoans. Current biology : CB. 25:1542–1550.

Desai, A., S. Rybina, T. Muller-Reichert, A. Shevchenko, A. Shevchenko, A. Hyman, and K. Oegema. 2003. KNL-1 directs assembly of the microtubule-binding interface of the kinetochore in C. elegans. Genes Dev. 17:2421–2435.

Dumont, J., and A. Desai. 2012. Acentrosomal spindle assembly and chromosome segregation during oocyte meiosis. Trends Cell Biol. 22:241–249.

Dumont, J., K. Oegema, and A. Desai. 2010. A kinetochore-independent mechanism drives anaphase chromosome separation during acentrosomal meiosis. Nat Cell Biol. 12:894–901.

El Yakoubi, W., and K. Wassmann. 2017. Meiotic Divisions: No Place for Gender Equality. Advances in experimental medicine and biology. 1002:1–17.

Enos, S.J., M. Dressler, B.F. Gomes, A.A. Hyman, and J.B. Woodruff. 2018. Phosphatase PP2A and microtubule-mediated pulling forces disassemble centrosomes during mitotic exit. Biol Open. 7.

Fabig, G., T. Muller-Reichert, and L.V. Paliulis. 2016. Back to the roots: segregation of univalent sex chromosomes in meiosis. Chromosoma. 125:277–286.

Fabig, G., A. Schwarz, C. Striese, M. Laue, and T. Müller-Reichert. 2019. *In situ* analysis of male meiosis in *C. elegans*. Methods Cell Biol. 152:in press.

Fabritius, A.S., M.L. Ellefson, and F.J. McNally. 2011. Nuclear and spindle positioning during oocyte meiosis. Curr Opin Cell Biol. 23:78–84.

Farhadifar, R., C.F. Baer, A.C. Valfort, E.C. Andersen, T. Muller-Reichert, M. Delattre, and D.J. Needleman. 2015. Scaling, Selection, and Evolutionary Dynamics of the Mitotic Spindle. Current biology : CB. 25:732–740.

Garcia-Mengual, E., J.C. Trivino, A. Saez-Cuevas, J. Bataller, M. Ruiz-Jorro, and X. Vendrell. 2019. Male infertility: establishing sperm aneuploidy thresholds in the laboratory. J Assist Reprod Genet. 36:371–381.

Ghongane, P., M. Kapanidou, A. Asghar, S. Elowe, and V.M. Bolanos-Garcia. 2014. The dynamic protein Knl1 - a kinetochore rendezvous. J Cell Sci. 127:3415–3423.

Grill, S.W., J. Howard, E. Schaffer, E.H. Stelzer, and A.A. Hyman. 2003. The distribution of active force generators controls mitotic spindle position. Science. 301:518–521.

Han, X., K. Adames, E.M. Sykes, and M. Srayko. 2015. The KLP-7 Residue S546 Is a Putative Aurora Kinase Site Required for Microtubule Regulation at the Centrosome in C. elegans. PloS one. 10:e0132593.

Hannak, E., M. Kirkham, A.A. Hyman, and K. Oegema. 2001. Aurora-A kinase is required for centrosome maturation in Caenorhabditis elegans. J Cell Biol. 155:1109–1116.

Hassold, T., and P. Hunt. 2001. To err (meiotically) is human: the genesis of human aneuploidy. Nat Rev Genet. 2:280–291.

Hauf, S., and Y. Watanabe. 2004. Kinetochore orientation in mitosis and meiosis. Cell. 119:317–327.

Hodgkin, J.A., and S. Brenner. 1977. Mutations causing transformation of sexual phenotype in the nematode Caenorhabditis elegans. Genetics. 86:275–287.

Howe, M., K.L. McDonald, D.G. Albertson, and B.J. Meyer. 2001. HIM-10 is required for kinetochore structure and function on Caenorhabditis elegans holocentric chromosomes. J Cell Biol. 153:1227–1238.

Ioannou, D., and H.G. Tempest. 2015. Meiotic Nondisjunction: Insights into the Origin and Significance of Aneuploidy in Human Spermatozoa. Advances in experimental medicine and biology. 868:1–21.

Kim, E., L. Sun, C.V. Gabel, and C. Fang-Yen. 2013. Long-term imaging of *Caenorhabditis elegans* using nanoparticle-mediated immobilization. PloS one. 8:e53419.

Kline, S.L., I.M. Cheeseman, T. Hori, T. Fukagawa, and A. Desai. 2006. The human Mis12 complex is required for kinetochore assembly and proper chromosome segregation. J Cell Biol. 173:9–17.

Kremer, J.R., D.N. Mastronarde, and J.R. McIntosh. 1996. Computer visualization of three-dimensional image data using IMOD. J Struct Biol. 116:71–76.

Kudalkar, E.M., E.A. Scarborough, N.T. Umbreit, A. Zelter, D.R. Gestaut, M. Riffle, R.S. Johnson, M.J. MacCoss, C.L. Asbury, and T.N. Davis. 2015. Regulation of outer kinetochore Ndc80 complex-based microtubule attachments by the central kinetochore Mis12/MIND complex. Proceedings of the National Academy of Sciences of the United States of America. 112:E5583–5589.

L’Hernault, S.W. 2006. Spermatogenesis. WormBook:1–14.

Laband, K., R. Le Borgne, F. Edwards, M. Stefanutti, J.C. Canman, J.M. Verbavatz, and J. Dumont. 2017. Chromosome segregation occurs by microtubule pushing in oocytes. Nat Commun. 8:1499.

LaFountain, J.R., Jr., C.S. Cohan, and R. Oldenbourg. 2011. Functional states of kinetochores revealed by laser microsurgery and fluorescent speckle microscopy. Mol Biol Cell. 22:4801–4808.

LaFountain, J.R., Jr., C.S. Cohan, and R. Oldenbourg. 2012. Pac-man motility of kinetochores unleashed by laser microsurgery. Mol Biol Cell. 23:3133–3142.

Levine, H., N. Jorgensen, A. Martino-Andrade, J. Mendiola, D. Weksler-Derri, I. Mindlis, R. Pinotti, and S.H. Swan. 2017. Temporal trends in sperm count: a systematic review and meta-regression analysis. Hum Reprod Update. 23:646–659.

Levine, H., H. Mohri, A. Ekbom, L. Ramos, G. Parker, E. Roldan, L. Jovine, S. Koelle, A. Lindstrand, S. Immler, S. Mortimer, D. Mortimer, G. van der Horst, S. Ishijima, N. Aneck-Hahn, E. Baldi, R. Menkveld, S.A. Rothmann, A. Giwercman, Y. Giwercman, M. Holmberg, U. Kvist, L. Bjorndahl, R. Holmberg, S. Arver, J. Flanagan, and J.R. Drevet. 2018. Male reproductive health statement (XIIIth international symposium on Spermatology, may 9th-12th 2018, Stockholm, Sweden. Basic Clin Androl. 28:13.

Lindow, N., S. Redemann, F. Brüning, G. Fabig, T. Müller-Reichert, and S. Prohaska. 2018. Quantification of three-dimensional spindle architecture. Methods Cell Biol. 145:45–64.

Ly, P., S.F. Brunner, O. Shoshani, D.H. Kim, W. Lan, T. Pyntikova, A.M. Flanagan, S. Behjati, D.C. Page, P.J. Campbell, and D.W. Cleveland. 2019. Chromosome segregation errors generate a diverse spectrum of simple and complex genomic rearrangements. Nat Genet. 51:705–715.

Maddox, P.S., K.D. Corbett, and A. Desai. 2012. Structure, assembly and reading of centromeric chromatin. Curr Opin Genet Dev. 22:139–147.

Madl, J.E., and R.K. Herman. 1979. Polyploids and sex determination in Caenorhabditis elegans. Genetics. 93:393–402.

Magescas, J., J.C. Zonka, and J.L. Feldman. 2019. A two-step mechanism for the inactivation of microtubule organizing center function at the centrosome. Elife. 8.

Mastronarde, D.N. 1997. Dual-axis tomography: an approach with alignment methods that preserve resolution. J Struct Biol. 120:343–352.

McIntosh, J.R. 2017. Mechanisms of Mitotic Chromosome Segregation. MDPI AG, Basel, Switzerland. McIntosh, J.R., M.I. Molodtsov, and F.I. Ataullakhanov. 2012. Biophysics of mitosis. Q Rev Biophys. 45:147–207.

McNally, K.P., M.T. Panzica, T. Kim, D.B. Cortes, and F.J. McNally. 2016. A novel chromosome segregation mechanism during female meiosis. Mol Biol Cell. 27:2576–2589.

Monen, J., P.S. Maddox, F. Hyndman, K. Oegema, and A. Desai. 2005. Differential role of CENP-A in the segregation of holocentric C. elegans chromosomes during meiosis and mitosis. Nat Cell Biol. 7:1248–1255.

Moore, L.L., and M.B. Roth. 2001. HCP-4, a CENP-C-like protein in Caenorhabditis elegans, is required for resolution of sister centromeres. J Cell Biol. 153:1199–1208.

Moore, L.L., G. Stanvitch, M.B. Roth, and D. Rosen. 2005. HCP-4/CENP-C promotes the prophase timing of centromere resolution by enabling the centromere association of HCP-6 in Caenorhabditis elegans. Mol Cell Biol. 25:2583–2592.

Muller-Reichert, T., G. Greenan, E. O’Toole, and M. Srayko. 2010. The *elegans* of spindle assembly. Cell Mol Life Sci. 67:2195–2213.

Muller-Reichert, T., H. Hohenberg, E.T. O’Toole, and K. McDonald. 2003. Cryoimmobilization and three-dimensional visualization of C. elegans ultrastructure. J Microsc. 212:71–80.

Muller-Reichert, T., J. Mantler, M. Srayko, and E. O’Toole. 2008. Electron microscopy of the early Caenorhabditis elegans embryo. J Microsc. 230:297–307.

Muscat, C.C., K.M. Torre-Santiago, M.V. Tran, J.A. Powers, and S.M. Wignall. 2015. Kinetochore-independent chromosome segregation driven by lateral microtubule bundles. Elife. 4:e06462.

Nahaboo, W., M. Zouak, P. Askjaer, and M. Delattre. 2015. Chromatids segregate without centrosomes during Caenorhabditis elegans mitosis in a Ran- and CLASP-dependent manner. Mol Biol Cell. 26:2020–2029.

Nicklas, R.B., and D.F. Kubai. 1985. Microtubules, chromosome movement, and reorientation after chromosomes are detached from the spindle by micromanipulation. Chromosoma. 92:313–324.

Nicklas, R.B., J.C. Waters, E.D. Salmon, and S.C. Ward. 2001. Checkpoint signals in grasshopper meiosis are sensitive to microtubule attachment, but tension is still essential. J Cell Sci. 114:4173–4183.

O’Donnell, L., and M.K. O’Bryan. 2014. Microtubules and spermatogenesis. Semin Cell Dev Biol. 30:45–54.

O’Toole, E.T., K.L. McDonald, J. Mantler, J.R. McIntosh, A.A. Hyman, and T. Muller-Reichert. 2003. Morphologically distinct microtubule ends in the mitotic centrosome of Caenorhabditis elegans. J Cell Biol. 163:451–456.

Oegema, K., A. Desai, S. Rybina, M. Kirkham, and A.A. Hyman. 2001. Functional analysis of kinetochore assembly in Caenorhabditis elegans. J Cell Biol. 153:1209–1226.

Peters, N., D.E. Perez, M.H. Song, Y. Liu, T. Muller-Reichert, C. Caron, K.J. Kemphues, and K.F. O’Connell. 2010. Control of mitotic and meiotic centriole duplication by the Plk4-related kinase ZYG-1. J Cell Sci. 123:795–805.

Petronczki, M., M.F. Siomos, and K. Nasmyth. 2003. Un menage a quatre: the molecular biology of chromosome segregation in meiosis. Cell. 112:423–440.

Petrovic, A., J. Keller, Y. Liu, K. Overlack, J. John, Y.N. Dimitrova, S. Jenni, S. van Gerwen, P. Stege, S. Wohlgemuth, P. Rombaut, F. Herzog, S.C. Harrison, I.R. Vetter, and A. Musacchio. 2016. Structure of the MIS12 Complex and Molecular Basis of Its Interaction with CENP-C at Human Kinetochores. Cell. 167:1028–1040 e1015.

Petrovic, A., S. Pasqualato, P. Dube, V. Krenn, S. Santaguida, D. Cittaro, S. Monzani, L. Massimiliano, J. Keller, A. Tarricone, A. Maiolica, H. Stark, and A. Musacchio. 2010. The MIS12 complex is a protein interaction hub for outer kinetochore assembly. J Cell Biol. 190:835–852.

Phillips, C.M., and A.F. Dernburg. 2006. A family of zinc-finger proteins is required for chromosome-specific pairing and synapsis during meiosis in C. elegans. Dev Cell. 11:817–829.

Phillips, C.M., C. Wong, N. Bhalla, P.M. Carlton, P. Weiser, P.M. Meneely, and A.F. Dernburg. 2005. HIM-8 binds to the X chromosome pairing center and mediates chromosome-specific meiotic synapsis. Cell. 123:1051–1063.

Pintard, L., and B. Bowerman. 2019. Mitotic Cell Division in Caenorhabditis elegans. Genetics. 211:35–73.

Reck-Peterson, S.L., W.B. Redwine, R.D. Vale, and A.P. Carter. 2018. The cytoplasmic dynein transport machinery and its many cargoes. Nat Rev Mol Cell Biol. 19:382–398.

Redemann, S., J. Baumgart, N. Lindow, M. Shelley, E. Nazockdast, A. Kratz, S. Prohaska, J. Brugues, S. Furthauer, and T. Muller-Reichert. 2017. C. elegans chromosomes connect to centrosomes by anchoring into the spindle network. Nat Commun. 8:15288.

Redemann, S., I. Lantzsch, N. Lindow, S. Prohaska, M. Srayko, and T. Muller-Reichert. 2018. A Switch in Microtubule Orientation during *C. elegans* Meiosis. Current biology : CB. 28:2991–2997.

Redemann, S., B. Weber, M. Moller, J.M. Verbavatz, A.A. Hyman, D. Baum, S. Prohaska, and T. Muller-Reichert. 2014. The segmentation of microtubules in electron tomograms using Amira. Methods Mol Biol. 1136:261–278.

Ris, H. 1949. The anaphase movement of chromosomes in the spermatocytes of the grasshopper. Biol Bull. 96:90–106.

Schindelin, J., I. Arganda-Carreras, E. Frise, V. Kaynig, M. Longair, T. Pietzsch, S. Preibisch, C. Rueden, S. Saalfeld, B. Schmid, J.Y. Tinevez, D.J. White, V. Hartenstein, K. Eliceiri, P. Tomancak, and A. Cardona. 2012. Fiji: an open-source platform for biological-image analysis. Nat Methods. 9:676–682.

Schmidt, D.J., D.J. Rose, W.M. Saxton, and S. Strome. 2005. Functional analysis of cytoplasmic dynein heavy chain in Caenorhabditis elegans with fast-acting temperature-sensitive mutations. Mol Biol Cell. 16:1200–1212.

Schmidt, R., L.E. Fielmich, I. Grigoriev, E.A. Katrukha, A. Akhmanova, and S. van den Heuvel. 2017. Two populations of cytoplasmic dynein contribute to spindle positioning in C. elegans embryos. J Cell Biol. 216:2777–2793.

Scholey, J.M., G. Civelekoglu-Scholey, and I. Brust-Mascher. 2016. Anaphase B. Biology (Basel). 5.

Schvarzstein, M., D. Pattabiraman, J.N. Bembenek, and A.M. Villeneuve. 2013. Meiotic HORMA domain proteins prevent untimely centriole disengagement during Caenorhabditis elegans spermatocyte meiosis. Proceedings of the National Academy of Sciences of the United States of America. 110:E898–907.

Sengupta, P., E. Borges, Jr., S. Dutta, and E. Krajewska-Kulak. 2018. Decline in sperm count in European men during the past 50 years. Hum Exp Toxicol. 37:247–255.

Severson, A.F., and B.J. Meyer. 2014. Divergent kleisin subunits of cohesin specify mechanisms to tether and release meiotic chromosomes. Elife. 3:e03467.

Severson, A.F., G. von Dassow, and B. Bowerman. 2016. Oocyte Meiotic Spindle Assembly and Function. Curr Top Dev Biol. 116:65–98.

Shakes, D.C., B.J. Neva, H. Huynh, J. Chaudhuri, and A. Pires-Dasilva. 2011. Asymmetric spermatocyte division as a mechanism for controlling sex ratios. Nat Commun. 2:157.

Shakes, D.C., J.C. Wu, P.L. Sadler, K. Laprade, L.L. Moore, A. Noritake, and D.S. Chu. 2009. Spermatogenesis-specific features of the meiotic program in Caenorhabditis elegans. PLoS genetics. 5:e1000611.

Skibbens, R.V., V.P. Skeen, and E.D. Salmon. 1993. Directional instability of kinetochore motility during chromosome congression and segregation in mitotic newt lung cells: a push-pull mechanism. J Cell Biol. 122:859–875.

Skop, A.R., D. Bergmann, W.A. Mohler, and J.G. White. 2001. Completion of cytokinesis in C. elegans requires a brefeldin A-sensitive membrane accumulation at the cleavage furrow apex. Current biology : CB. 11:735–746.

Soppina, V., A.K. Rai, A.J. Ramaiya, P. Barak, and R. Mallik. 2009. Tug-of-war between dissimilar teams of microtubule motors regulates transport and fission of endosomes. Proceedings of the National Academy of Sciences of the United States of America. 106:19381–19386.

Srayko, M., T. O’Toole E, A.A. Hyman, and T. Muller-Reichert. 2006. Katanin disrupts the microtubule lattice and increases polymer number in C. elegans meiosis. Current biology : CB. 16:1944–1949.

Stalling, D., M. Westerhoff, and H.-C. Hege. 2005. Amira: a highly interactive system for visual data analysis. In The Visualization Handbook. C.D. Hansen and C.R. Johnson, editors. Elsevier. 749–767.

Strome, S., J. Powers, M. Dunn, K. Reese, C.J. Malone, J. White, G. Seydoux, and W. Saxton. 2001. Spindle dynamics and the role of gamma-tubulin in early Caenorhabditis elegans embryos. Mol Biol Cell. 12:1751–1764.

Sulston, J., and J. Hodgkin. 1988. Methods. *In* The nematode C. elegans. B.W. Wood, editor. Cold Spring Harbor Laboratory Press, Cold Spring Harbor, New York. 587–606.

Sutradhar, S., and R. Paul. 2014. Tug-of-war between opposing molecular motors explains chromosomal oscillation during mitosis. J Theor Biol. 344:56–69.

Weber, B., G. Greenan, S. Prohaska, D. Baum, H.-C. Hege, T. Müller-Reichert, A.A. Hyman, and J.-M. Verbavatz. 2012. Automated tracing of microtubules in electron tomograms of plastic embedded samples of *C. elegans* embryos. J Struct Biol. 178:129–138.

Weber, B., E.M. Tranfield, J.L. Hoog, D. Baum, C. Antony, T. Hyman, J.M. Verbavatz, and S. Prohaska. 2014. Automated stitching of microtubule centerlines across serial electron tomograms. PloS one. 9:e113222.

Wei, R.R., J. Al-Bassam, and S.C. Harrison. 2007. The Ndc80/HEC1 complex is a contact point for kinetochore-microtubule attachment. Nat Struct Mol Biol. 14:54–59.

Wilson-Kubalek, E.M., I.M. Cheeseman, and R.A. Milligan. 2016. Structural comparison of the Caenorhabditis elegans and human Ndc80 complexes bound to microtubules reveals distinct binding behavior. Mol Biol Cell. 27:1197–1203.

Winter, E.S., A. Schwarz, G. Fabig, J.L. Feldman, A. Pires-daSilva, T. Muller-Reichert, P.L. Sadler, and D.C. Shakes. 2017. Cytoskeletal variations in an asymmetric cell division support diversity in nematode sperm size and sex ratios. Development. 144:3253–3263.

Yu, C.-H., S. Redemann, H.-Y. Wu, R. Kiewisz, T.Y. Yoo, R. Farhadifar, T. Müller-Reichert, and D. Needleman. 2019. Central spindle microtubules are strongly coupled to chromosomes during both anaphase A and anaphase B. Mol Biol Cell. doi: 10.1091/mbc.E19-01-0074.

Zhang, D., and R.B. Nicklas. 1995. Chromosomes initiate spindle assembly upon experimental dissolution of the nuclear envelope in grasshopper spermatocytes. J Cell Biol. 131:1125–1131.

